# Structural evidence for MADS-box type I family expansion seen in new assemblies of *A. arenosa* and *A. lyrata*

**DOI:** 10.1101/2023.05.30.542816

**Authors:** Jonathan Bramsiepe, Anders K. Krabberød, Katrine N. Bjerkan, Renate M. Alling, Ida M. Johannessen, Karina S. Hornslien, Jason R. Miller, Anne K. Brysting, Paul E. Grini

## Abstract

*Arabidopsis thaliana* diverged from *A. arenosa* and *A. lyrata* at least 6 million years ago and are identified by genome-wide polymorphisms or morphological traits. The species are to a high degree reproductively isolated, but hybridization barriers are incomplete. A special type of hybridization barrier is based in the triploid endosperm of the seed, where embryo lethality is caused by endosperm failure to support the developing embryo. The MADS-box type I family of transcription factors are specifically expressed in the endosperm and has been proposed to play a role in endosperm-based hybridization barriers. The gene family is well known for a high evolutionary duplication rate, as well as being regulated by genomic imprinting. Here we address MADS-box type I gene family evolution and the role of type I genes in the context of hybridization. Using two *de-novo* assembled and annotated chromosome-level genomes of *A. arenosa* and *A. lyrata* ssp. *petraea* we analyzed the MADS-box type I gene family in *Arabidopsis* to predict orthologs, copy number and structural genomic variation related to the type I loci. Our findings were compared to gene expression profiles sampled before and after the transition to endosperm cellularization in order to investigate the involvement of MADS-box type I loci in endosperm-based hybridization barriers. We observed substantial differences in type-I expression between *A. arenosa* and *A. lyrata* ssp. *petraea* in the endosperm, suggesting a genetic cause for the endosperm-based hybridization barrier in *A. arenosa* and *A. lyrata* ssp. *petraea* hybrid seeds.

## Introduction

*Arabidopsis thaliana* diverged from its closest relative, *A. arenosa* and *A. lyrata*, at least 6 million years ago (mya) (Hohmann *et al*., 2015), corresponding with the basal chromosome number reduction from eight to five in *A. thaliana (Lysak et al., 2006)*. The species are identified by genome-wide polymorphisms or morphological traits, but only the monophyly of *A. thaliana* is convincingly described as being supported 100% at the level of individual gene trees (Novikova *et al*., 2016). The species are to a high degree reproductively isolated, but natural field studies (Schmickl and Koch, 2011; Marburger *et al*., 2019; Schmickl and Yant, 2021) and interspecific hybridization in controlled conditions (Chen *et al*., 1998; Comai *et al*., 2000; Nasrallah *et al*., 2000; Josefsson *et al*., 2006; Walia *et al*., 2009; Burkart-Waco *et al*., 2012; Burkart-Waco *et al*., 2013; Burkart-Waco *et al*., 2015; Bjerkan *et al*., 2020) show that hybridization barriers are incomplete.

A special type of hybridization barrier is based in the triploid endosperm of the seed, where a syncytial growth phase accumulating storage components followed by cellularization switches the endosperm from nutrition sink to the primary nutrition source of the embryo (Hehenberger *et al*., 2012). Embryo lethality is frequently observed in hybrid seeds and contrasted by embryo survival when cultivated *in vitro* after microdissection, suggesting that the endosperm fails to support and transfer nutrition to the embryo (Rebernig *et al*., 2015; Florez-Rueda *et al*., 2016; Tonosaki *et al*., 2017; Lafon-Placette *et al*., 2017).

The MADS-box type I family of transcription factors are specifically expressed in the endosperm at the time of transition to cellularization (Shirzadi *et al*., 2011; Masiero *et al*., 2011; Bjerkan *et al*., 2020; Bemer *et al*., 2010; Zhang *et al*., 2018), and several lines of evidence has suggested a role of type I genes in endosperm based hybridization barriers (Josefsson *et al*., 2006; Walia *et al*., 2009; Bjerkan *et al*., 2020; Burkart-Waco *et al*., 2015). The gene family is well known for its high duplication rate, and is contrasted by its sister lineage, the MADS-box type II family, which consists of highly conserved single-copy genes (Parenicová *et al*., 2003; Gramzow and Theissen, 2010; Qiu and Köhler, 2022). Members of the MADS-box type I family are also well known for being regulated by genomic imprinting, parent of origin dependent allelic expression in the endosperm (Köhler *et al*., 2003; Shirzadi *et al*., 2011; Bjerkan *et al*., 2020; Zhang *et al*., 2018; Masiero *et al*., 2011).

Whether gene duplication in the MADS-box type I family is favored or sensed by genomic imprinting is disputed (Nam *et al*., 2004; Yoshida and Kawabe, 2013; Erilova *et al*., 2009). One of the first discovered imprinted genes, *MEDEA* (*MEA*), is crucial for endosperm cellularization (Grossniklaus *et al*., 1998; Kinoshita *et al*., 1999), and a primary target of MEA is *PHERES1* (*PHE1*), a paternally expressed MADS-box type I transcription factor. A reduction of *PHE1* expression rescues the *mea* mutant phenotype in seeds, and in hybrid crosses, parental imprinting of *PHE1* is altered to the opposite parent (Köhler *et al*., 2003; Josefsson *et al*., 2006; Walia *et al*., 2009). Recently, a propagation of PHERES1 target sites by transposon events was described and indicated a powerful and recent evolutionary effect (Batista *et al*., 2019; Qiu and Köhler, 2020). Addressing the involvement of MADS-box type I genes and genomic imprinting in the context of hybridization, and especially in crosses between outcrossing species, is important since the occurrence of the phenomenon in selfing species may be considered an evolutionary remnant from outcrossing species.

Such analyses require well-assembled genomes to allow for correct ortholog prediction and identification of paralogs, and the ability to address structural variation in the DNA sequences. The *Arabidopsis thaliana* genome caused an explosion in plant genome research, supporting questions in many fields (The Arabidopsis Genome Initiative, 2000). The small genome size and the strong homozygosity of *A. thaliana* made this effort successful but is not archetypical for the majority of plants, not even the genus *Arabidopsis*. The complexity of whole-genome duplications, ancient and recent polyploidization events, and also high transposon activity creating abundant pseudogenes, make the assembly of plant genomes especially challenging (Kress *et al*., 2022). Thus, for many sequenced plant genomes, the assembly of repeats such as centromeres, telomeres, and ribosomal DNA has become feasible only after considerable methodological improvements implemented by long-read sequencing technology and Hi-C scaffolding (Naish *et al*., 2021; Kovaka *et al*., 2023).

Using assembled *Arabidopsis* genomes and new assemblies of the genomes of the European *A. arenosa* and *A. lyrata* ssp. *petraea*, we have analyzed the MADS-box type I gene family in *Arabidopsis* to predict orthologs, copy number and structural genomic variation related to the type I loci. The analysis is compared to gene expression profiles sampled before and after the transition to endosperm cellularization in order to investigate the involvement of MADS-box type I loci in endosperm based hybridization barriers.

## Results and Discussion

### Assembly and annotation of *A. arenosa* and *A. lyrata* ssp. *petraea* genomes

To identify MADS-box type I family orthologs between species, allowing interspecies gene expression studies of orthologs between *Arabidopsis* species, and comparative analysis of orthologous and paralogous genes including their genomic neighborhood, we sequenced and assembled the genomes of one individual of *A. arenosa* from Pusté Pole in Slovakia and one individual of *A. lyrata* ssp. *petraea* from Pernitz in Austria (see Figure S1 and Data S1 for a summary of assembly methods and statistics). The genome sizes were estimated as 201 Mbp for *A. lyrata* ssp. *petraea* and 179 Mbp for *A. arenosa* Pusté Pole (see K-mer analysis in Figure S2), which were confirmed by flow-cytometric data (Data S2) and previous reports (Johnston *et al*., 2005; Lysak *et al*., 2009; Dart *et al*., 2004), and by the assembled sequence lengths of 188 Mbp and 153 Mbp. Whole-genome alignments between scaffolded assemblies were analyzed for breaks and inconsistencies (Figure S3-6(Robinson *et al*., 2011; Durand *et al*., 2016; Dudchenko *et al*., 2018), revealing chromosome-length scaffolds comparable in quality to other published genomes of *Arabidopsis* (Data S1).

Localization of rDNA, telomeric and centromeric repeats indicate complete assemblies and the remaining gaps clustered close to the centromeres (Figures 1A and B). Repeats and transposable elements (TE) were classified and compared to their occurrence in *A. lyrata* ssp. *lyrata* and *A. thaliana* genomes, revealing a higher percentage of repeat sequences in *A. arenosa* and *A. lyrata* compared to *A. thaliana* (Figure 1C). Comparison of the predicted proteomes of *A. arenosa* Pusté Pole, *A. lyrata* ssp. *petraea,* and *A. thaliana* detected more than 21 thousand shared orthogroups and a few (<2%) species-specific orthogroups (Figure 1D). Orthogroups shared by the *A. arenosa* and *A. lyrata* ssp. *petraea* species only were two to three times more frequent, reflecting the recent split of the two species. Analysis of local structural variants of *A. arenosa* Pusté Pole and *A. lyrata* ssp. *petraea* to *A. thaliana* and *A. lyrata* ssp. *lyrata* revealed a high syntenic relationship and limited chromosomal rearrangement (Figure 1E, Figure S7).

**Figure 1.**
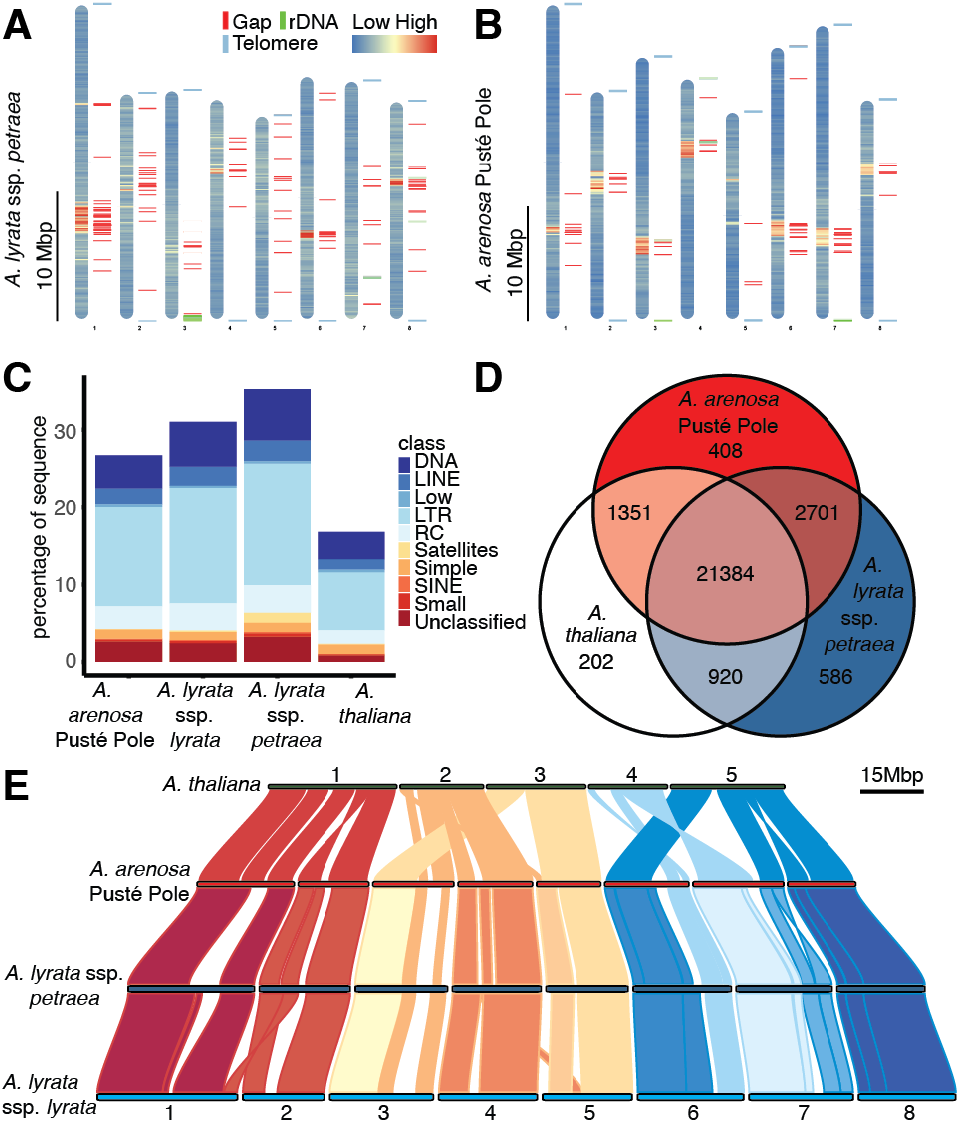
Comparison of *Arabidopsis* genome assemblies. Repeats and genes were predicted using RepeatModeler and Braker. (A-B) Idiograms of *A. lyrata* ssp. *petraea* (A) and *A. arenosa* Pusté Pole (B). The repeat density is shown as a heat map. Gaps (red), rDNA (green), and telomeric sequences (blue) are indicated as bars next to the chromosomes. Hi-C scaffolding gives a continuous genome assembly, and the few remaining gaps cluster in centromeric regions. (C) Abundance of repeat classes in *A. lyrata* ssp. *petraea* and *A. arenosa* Pusté Pole compared to *A. thaliana* and *A. lyrata* ssp. *lyrata*. The *A. thaliana* genome has the lowest number of repeats relative to genome size. The increased number of repetitive elements in *A. lyrata* ssp*. petraea* compared to *A. lyrata* ssp. *lyrata* is probably due to the technical advantage of using long read sequencing technology (Pucker *et al*., 2022). (D) Venn diagram showing overlap of predicted orthologs between *A. lyrata* ssp. *petraea, A. arenosa* Pusté Pole, and *A. thaliana*. Orthogroups were predicted using OrthoFinder. A majority of orthologs are common for all three species, while few species-specific genes were identified. (E) Synteny of *A. lyrata* ssp. *petraea* and *A. arenosa* Pusté Pole assemblies to *A. lyrata* ssp. *lyrata* and *A. thaliana* references. Known rearrangements can be found in the comparison to the n=5 based karyotype in *A. thaliana*. The karyotypes between *A. arenosa* and *A. lyrata* overlap, with one large inversion on the end of scaffold 7. The *A. lyrata* ssp. *petraea* assembly was corrected for a previously reported misassembly on scaffold 1 (Slotte *et al*., 2013).

### MADS-box gene family characterization identifies diagnostic motifs

To identify MADS-box type I family orthologs between species, gene sequences from *A. thaliana* were used to identify the MADS-box genes in four currently available *Arabidopsis* genomes of sufficient quality (*A. thaliana, A. lyrata* ssp. *lyrata*, *A. halleri*, and *A. arenosa* Strecno) in addition to the two newly assembled genomes (see Data S3.1 for all sequences). In a phylogenetic tree using two *Capsella* species as outgroups (*C. rubella* and *C. grandiflora*), the MADS-box genes were categorized into types and subgroups according to phylogenetic placement (Figure 2, Figure S8, Figure S9).

**Figure 2.**
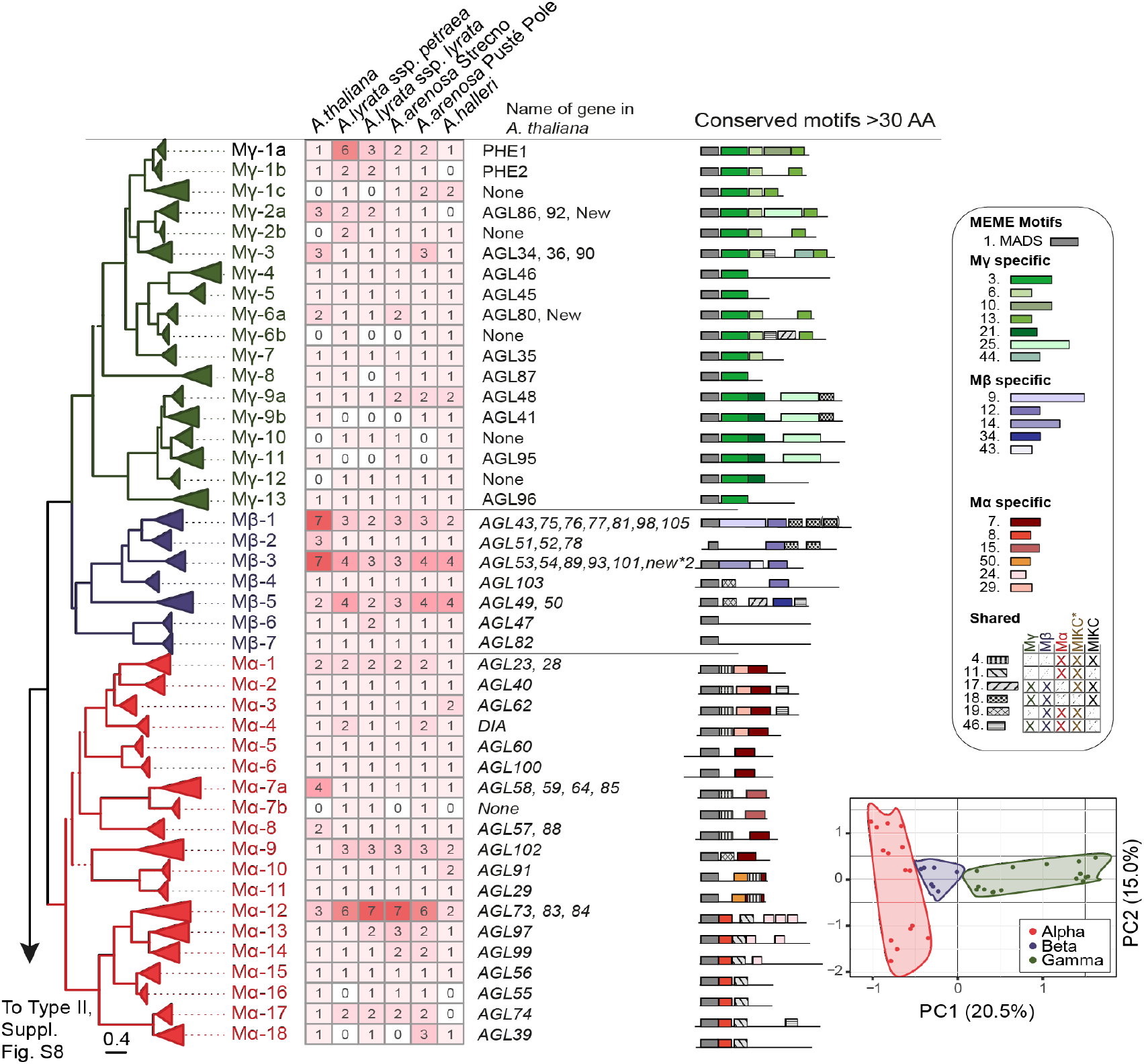
Phylogenetic analysis of MADS-box type I genes in *Arabidopsis*. The tree was derived by a maximum likelihood analysis of 362 identified MADS-box type I sequences from *A. thaliana, A. lyrata* ssp. *petraea, A. arenosa* Strecno*, A. arenosa* Pusté Pole, and *A. halleri* with two species of *Capsella* (100 sequences; not shown in the figure) used as outgroup. Solid branches represent bootstrap support > 85%, while branches with support values < 85% are dashed. The root of the tree is placed between type I and type II genes (the corresponding tree and heatmap for type II genes can be found in Figure S9). Triangles represent clades where branches are collapsed at the most recent gene duplication event in the last common ancestor of the genus *Arabidopsis*. The length of the triangles corresponds to the overall branch length of the collapsed clade. The groups are colored with Mα in red, Mβ in blue, and Mγ in green (see the main text for naming schemes of clades). The heatmap shows the number of gene copies for each clade in the five genomes. The column next to the heatmap indicates canonical MADS-box names of the genes in *A. thaliana* found in the corresponding clade; “none” means that a gene representing the clade is not found in the *A. thaliana* genome, while “new” indicates that the gene does not have a given MADS-box name in *A. thaliana*. The corresponding *A. thaliana* gene locus code (AGI) and commonly used synonymous names for MADS-box genes can be found in Data S3. The last column shows a simplified representation of the MEME motifs. The insert in the lower right corner shows PCA on the occurrence of MEME motifs (Full PCA in Figure S10). A fully expanded phylogenetic tree with individual tip labels, support values for all branches, and the outgroup *Capsella* can be found in Figure S8. Results from the full MEME analysis can be found in Figure S8.

As expected, the resulting tree separates type I from type II, as well as the 3 subgroups of type I (Mα, Mβ, Mγ) and the 2 subgroups of type II (MIKC and MIKC* [also referred to as Mδ]) (Henschel *et al*., 2002; Parenicová *et al*., 2003; Arora *et al*., 2007; Thangavel and Nayar, 2018; Gramzow and Theißen, 2013; Qiu and Köhler, 2022). Monophyletic groups of *Arabidopsis* MADS-box genes, sharing a common ancestor with genes from the outgroup *Capsella,* are numbered starting from the top of the tree. Additional gene duplications that occurred in the common ancestor of *Arabidopsis*, but after *Arabidopsis* and *Capsella* separated, are marked with an additional letter. For instance, “Mγ-1a” and “Mγ-1b” indicate that these two clades share a last common ancestor with *C. rubella* (i.e. “Mγ-1”), but have a gene duplication specific for *Arabidopsis* that occurred after the separation of *Arabidopsis* from the rest of the Brassicales (Figure 2; for a fully expanded tree with names of all groups and sequences, see Figure S8, Data S3).

A principal component analysis of MEME motifs showed that the distribution of sequence motifs follows the same patterns as the phylogenetic tree (Figure 2, Figure S10). This allowed the identification of motifs that are diagnostic for clades or even larger groups, as exemplified by Mγ motif 3 or Mα motifs 7 and 8 (Figure 2). The protein motifs in MADS-box genes were generally found to be more similar inside clades than between clades. Some motifs were distributed in all clades except one, as exemplified by motif 46 (not in MIKC) and motif 17 (not in Mα). Motifs that classified MADS-box type I gene sequences into one of the Mα, Mβ, and Mγ clades are shown in Figure 2, Figures S8 and S9.

### Variance in MADS-box type I gene copy number

The gene copy number was determined for each *Arabidopsis* species in each clade of the phylogenetic tree (Figure 2). The gene copy number varies for the MADS-box type I gene clades. The number of copies is low for all clades in type II MIKC (Figure S9) except the type II clade *FLOWERING LOCUS C* (*FLC*), which spans MIKC28 to MIKC32, including *MAF1*-*MAF5* and *FLC* (Figure S9), and which is arranged as a tandem array on chromosome 5 in *A. thaliana* (Ratcliffe *et al*., 2003). The *FLC* duplication in *A. lyrata* ssp. *petraea* and its effect on flowering were previously studied (Kemi *et al*., 2013).

Members of the Mβ group were highly duplicated. The Mβ-1, Mβ-3, and Mβ-5 clades have three to four copies in *A. arenosa* and *A. lyrata. A. thaliana* has seven copies in the Mβ-1 and Mβ-3 clades and two copies in the Mβ-5 clade (Figure 2). The Mα group is separated into two monophyletic groups (Parenicová *et al*., 2003). In order to investigate the biological roles of the two groups, we compared the newly assembled versions of those genes to previously published interaction maps between MADS-box proteins (Folter *et al*., 2005; Qiu and Köhler, 2022). The first group (Mα-1 to Mα-11) contains clades with proteins for which only interaction with Mγ proteins has been found. Mα-5 and Mα-9 are the exceptions for which protein interaction with the Mβ group was also found in a yeast two-hybrid screen (Folter *et al*., 2005; Qiu and Köhler, 2022). In strong contrast, for the second group (Mα-12 to Mα-18), all *A. thaliana* proteins could interact with proteins of the Mβ class, while Mα-13, Mα-14, Mα-17, and Mα-19 could also interact with Mγ in the same yeast two-hybrid studies (Qiu and Köhler, 2022; Folter *et al*., 2005).

The groups are also distinguished by gene copy number, with the second group having higher numbers comparable to the Mβ class. The Mα-1 (*AGL23/28*) and Mα-4 (*DIANA*) clades, which are involved in female gametophyte development in *A. thaliana* (Colombo *et al*., 2008; Bemer *et al*., 2008; Steffen *et al*., 2008), are each duplicated once in *A. lyrata* and *A. arenosa* Pusté Pole. The Mα-2 (*AGL40*) and the Mα-3 clade containing *AGL62* in *A. thaliana* are single-copy genes in *A. thaliana*, *A. lyrata*, and *A. arenosa*. A possible explanation for the lack of expansion in Mα-2 and Mα-3 is their functional requirement in endosperm development. Mutations in *AGL62* cause early endosperm cellularization in *A. thaliana,* and it is suggested that *AGL40* may have functions in the same pathway since *agl40* mutant plants produce smaller seeds (Kang *et al*., 2008; Roszak and Köhler, 2011; Kirkbride *et al*., 2019). Like Mβ-1 and Mβ-3, the Mα-7a clade is notable for its high duplication rate in *A. thaliana*: four duplicate genes (*AGL58*, *AGL59*, *AGL64* and *AGL85*) were found in the Mα-7a clade, while only single orthologs were detected in *A. lyrata* and *A. arenosa*. The opposite is found for the Mα-9 clade, which contains three loci in *A. lyrata* and *A. arenosa* but only one single gene in *A. thaliana* (*AGL102*). In the second Mβ-binding group, the Mα-12 stands out with high duplication rates in *A. lyrata* and *A. arenosa*, represented by *AGL73*, *AGL83*, and *AGL84* in *A. thaliana*. The other Mβ-binding Mα clades have, except for Mα-15, single-gene duplications or losses depending on the *Arabidopsis* species.

In the Mγ class, the Mγ-1 and Mγ-3 clades have variable MADS-box type I gene copy numbers. Mγ-1a, known as *PHERES1* (*PHE1*) in *A. thaliana*, shows six copies in *A. lyrata* ssp. *petraea* but only three in *A. lyrata* ssp. *lyrata*. The homolog *PHE2* in *A. thaliana* is grouped in the distinct Mγ-1b clade and duplicated in both *A. lyrata* subspecies. The Mγ-3 clade has three copies, *AGL34*, *AGL36*, and *AGL90* in *A. thaliana*, and a triplication can also be found in *A. arenosa* Pusté Pole contrasted by single-copy loci in the other species studied (Figure 2).

### Distribution and clustering of MADS-box type I genes differ in *Arabidopsis*

We compared gene family distributions on chromosomes between *A. thaliana* and other species in the genus. Whereas *A. thaliana,* MADS-box type II genes are highly conserved but distributed across all chromosomes (Parenicová *et al*., 2003; Gramzow and Theissen, 2010; Qiu and Köhler, 2022), the type I genes are primarily localized on chromosomes 1 and 5, hypothesized to be a result of recent and local duplications (Parenicová *et al*., 2003). In *A. lyrata* and *A. arenosa*, however, we find that the MADS-box type I genes are distributed across all eight chromosomes (Figure 3). The highly variable type I clades cluster in the vicinity of centromeric repeats.

**Figure 3.**
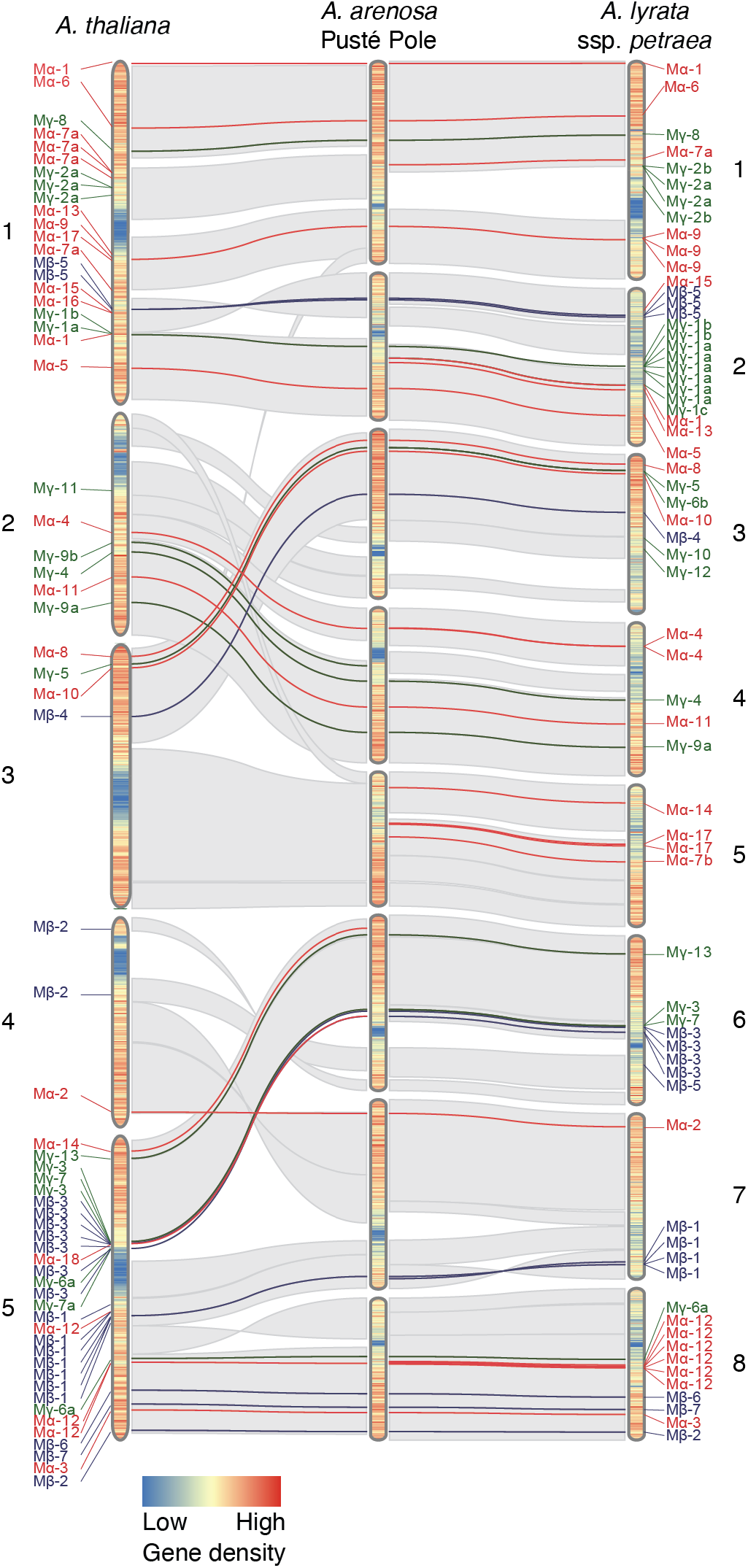
Genome-wide distribution of MADS-box type I genes in *Arabidopsis arenosa* and *A. lyrata* ssp. *petraea*. Localization of MADS-box type I Mα (red), Mβ (blue), and Mγ (green) are shown on top of the predicted syntenic sequence context. The naming corresponds to clades in the MADS-box phylogeny in Figure 2. While the MADS-box type I genes are mainly located on chromosomes 1 and 5 in *A. thaliana*, they are broadly distributed across all chromosomes of *A. arenosa* Pusté Pole and *A. lyrata* ssp. *petraea*. Local duplication and clusters of paralogs can be found in all three species. The heatmap covering the idiogram indicates gene density. Scaffolds are not in scale comparing species but show the proportion of the individual genomes (for scale, see Figure 1E and Figure S7).

Chromosomal localisation may explain the high duplication rate of type I clades in *A. lyrata* and *A. arenosa* compared to *A. thaliana*. For example, the variable Mγ-2 group is found close to centromeric repeats in *A. thaliana*, *A. lyrata*, and *A. arenosa*. There are three Mγ-2 in *A. thaliana*, four in *A. lyrata* ssp*. petraea*, and two in *A. arenosa* Pusté Pole -all on chromosome 1 (Figure 3). On the other hand, the Mγ-1 clades with *PHE1* and *PHE2* are located distantly from the centromere on the lower arm of chromosome 1 in *A. thaliana* and contain only these two genes. In *A. lyrata* ssp. *petraea* and *A. arenosa* Pusté Pole, the local duplications in these Mγ-1 clades are clustered close to the centromere of chromosome 2.

A similar pattern can be seen in regard to *A. thaliana* chromosome 5. The highly variable Mβ-3 cluster is located close to the centromeric repeats on chromosome 5 of *A. thaliana* and on chromosome 6 of *A. lyrata* ssp. *petraea* and *A. arenosa* Pusté Pole. In contrast, the Mα-12 loci are centrally located on the lower arm of chromosome 5 of *A. thaliana*, distant from the centromere, yet have a syntenic relationship to a region close to the centromeric repeats on chromosome 8 of *A. lyrata* ssp. *petraea* and *A. arenosa* Pusté Pole (Figure 3).

*AGL23* and *AGL28* of the Mα-1 clade are located on alternate chromosome arms of chromosome 1 of *A. thaliana*. The distant localization is maintained in syntenic regions in *A. lyrata* and *A. arenosa*, transferred to chromosomes 1 and 2 (Figure 3). The separate localization of these genes is consistent with their distinct expression (Figure 4). Likewise, *DIANA* (*AGL61*/Mα-4) is a single-copy gene on chromosome 2 of *A. thaliana*. However, the Mα-4 clade has a local duplication on chromosome 4 of *A. lyrata* and *A. arenosa* (Figure 3). Also, these genes show distinct expression during seed development (Figure 4).

**Figure 4.**
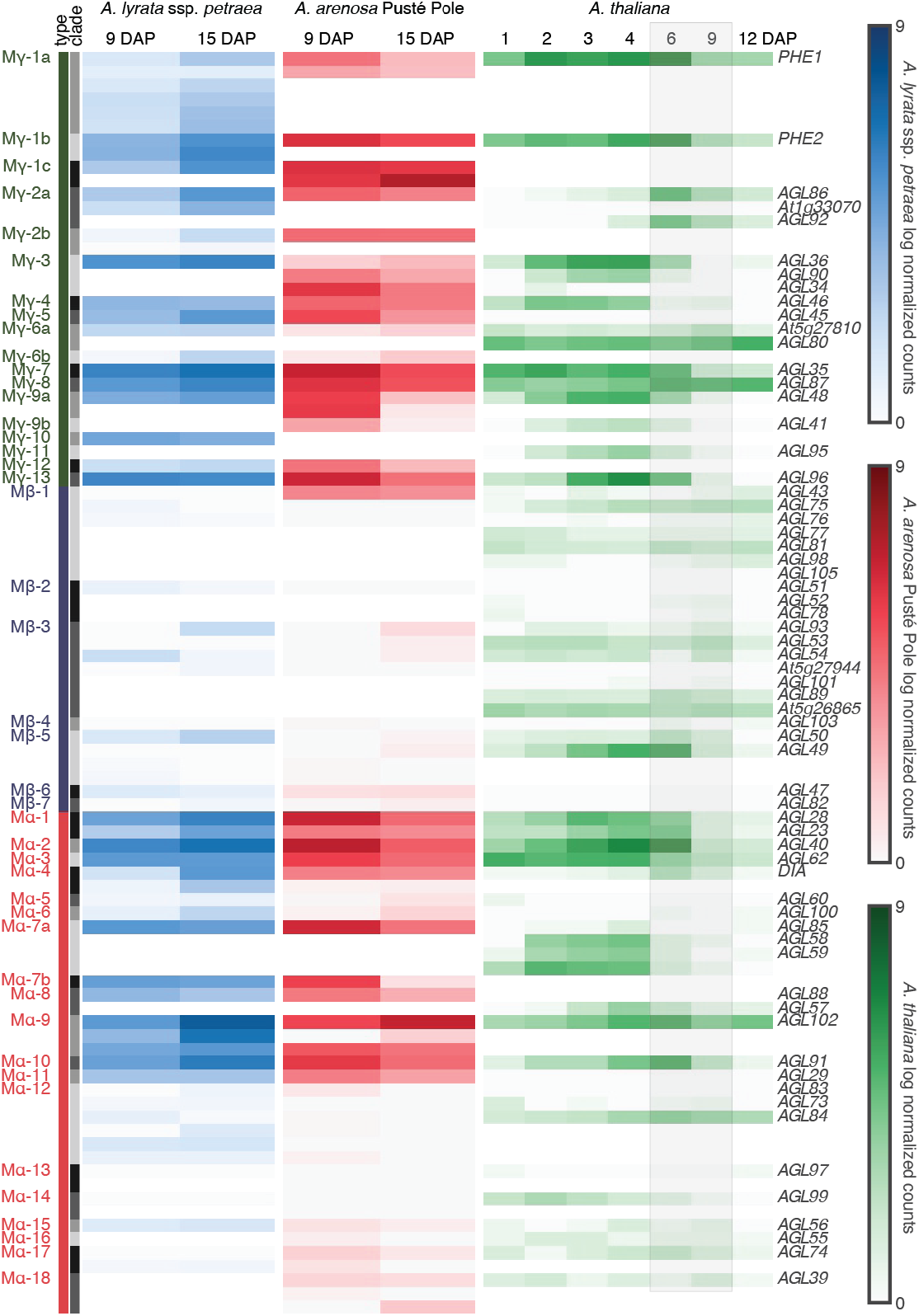
MADS-box type I expression during *Arabidopsis* seed development. Gene expression profiles are displayed for all identified MADS-box type I genes and compared between *A. lyrata* ssp. *petraea* (left column), *A. arenosa* Pusté Pole (middle column), and *A. thaliana* (right column). The *A. thaliana* development time series serves as a reference (Bjerkan *et al*., 2020). The endosperm cellularization in *A. thaliana* occurs between 6 and 9 days after pollination (DAP), indicated by the gray boxed area. To adjust for the relatively slower development in *A. arenosa* and *A. lyrata*, corresponding stages before and after endosperm cellularization were sampled at 9 and 15 DAP, respectively. Orthologous genes are grouped and ordered and follow the MADS-box phylogeny described in Figure 2. Counts normalized for each sample are shown on a base-2 logarithmic scale. Most Mγ and half of the Mα genes show a high expression before cellularization and a substantial decline post cellularization in *A. thaliana* and *A. arenosa*. In *A. lyrata* ssp. *petraea*, however, this decline of expression can not be found; instead, most of the expressed Mγ and Mα genes even increase in expression at the later (15 DAP) stage.

### Differential expression of MADS-box orthologs in *Arabidopsis* seeds

In order to further investigate the relationship between MADS-box type I expansion, their genomic neighborhood and gene expression in the seed, we generated seed transcript profiles of *A. lyrata* ssp. *petraea* and *A. arenosa* Pusté Pole. Several MADS-box type I genes regulate early seed development in *A. thaliana* and are expressed in an imprinted or biparental manner at the transition to endosperm cellularization (Bemer *et al*., 2010; Zhang *et al*., 2018; Bjerkan *et al*., 2020; Shirzadi *et al*., 2011; Masiero *et al*., 2011). In *A. thaliana,* a steep decline in MADS-box type I gene expression can be seen between 6 DAP and 9 DAP (days after pollination), coinciding with developmental stages before and after cellularization (Bjerkan *et al*., 2020) (Figure 4). Seed development in *A. lyrata* and *A. arenosa* progresses slower, and the comparable developmental phenotypes can be staged to 9 and 15 DAP (Lafon-Placette *et al*., 2017). We therefore compared MADS-box type I RNA-Seq profiles from 9 and 15 DAP *A. lyrata* ssp. *petraea* and *A. arenosa* Pusté Pole seeds to *A. thaliana* 6 to 9 DAP seed transcriptomes. While individual MADS-box type II genes were either never expressed or constantly expressed at both sampled stages in *A. thaliana* (Figure S11), their expression levels differed in *A. lyrata* ssp. *petraea* and *A. arenosa* Pusté Pole (Figure 4). In all species, high expression of the majority of Mγ genes was observed before cellularization, whereas Mβ expression is low or absent, and undergoes only minor changes during later stage endosperm development.

The Mα group expression pattern reflects its phylogeny and splits into two groups (Figure 2), as only the Mα-1 to Mα-11 clades are significantly expressed during early seed development. The weakly expressed Mα genes and the whole group of Mβ were characterized by high duplication levels. Although not significant for comparing all type-I MADS genes (R = -0.15 p = 0.077) we suggest that recent gene duplication in MADS-box type I genes correlates negatively with the observed seed-specific expression of these loci.

The Mα and Mγ expression patterns also reveal differences between *A. lyrata* ssp. *petraea* and *A. arenosa* Pusté Pole. In *A. arenosa*, the expression patterns are similar to *A. thaliana* with an up and down-regulation before and after endosperm cellularization, respectively. Surprisingly this pattern can not be found in *A. lyrata* ssp. *petraea*, where the expression of most MADS-box type I genes continues to increase at the later time point, sampled after endosperm cellularization. Mα-3 (*AGL62*), of which mutation leads to precocious cellularization in *A. thaliana* (Kang *et al*., 2008), is down-regulated at cellularization in *A. thaliana* and *A. arenosa* but its expression remains unchanged in *A. lyrata* ssp. *petraea.* Likewise, *Mγ-1a* (*PHE1*) expression declines in *A. thaliana* with the onset of endosperm cellularization, and *Mγ-1a* is also higher expressed during early seed development in *A. arenosa* and declines at cellularization. In *A. lyrata* ssp. *petraea,* however the *Mγ-1a* duplications show a higher level of expression at the 15 DAP time point compared to 9 DAP (Figure 4). In summary, the observed down-regulation of type I class Mα and Mγ in *A. thaliana* and *A. arenosa* is absent, weakened, or strongly delayed in *A. lyrata* ssp. *petraea*, or that endosperm cellularization occurs independent of MADS-box type I expression.

### Recent duplication in Mγ 1 (*PHERES1*)

The complete *A. lyrata* ssp. *petraea* and *A. arenosa* Pusté Pole genomes allow interspecies comparisons of gene family expansion. We investigated the observed lack of *Mγ-1a* down-regulation and extensive expansion of this clade in *A. lyrata* ssp. *petraea.* In *A. thaliana*, *Mγ-1a* (*PHE1*) is paternally expressed while the maternal allele is repressed by the POLYCOMB REPRESSIVE COMPLEX 2 (PRC2). It is proposed that PRC2 repression is circumvented in the paternal allele of *PHE1* by DNA-methylation of a repeat-rich region in the three-prime regulatory region of *PHE1 (Makarevich et al., 2008)*. We therefore used the whole-genome assemblies of *A. lyrata* ssp. *petraea* and *A. arenosa* Pusté Pole to address and compare the Mγ-1 duplications in their genomic landscape (Figure 5).

**Figure 5.**
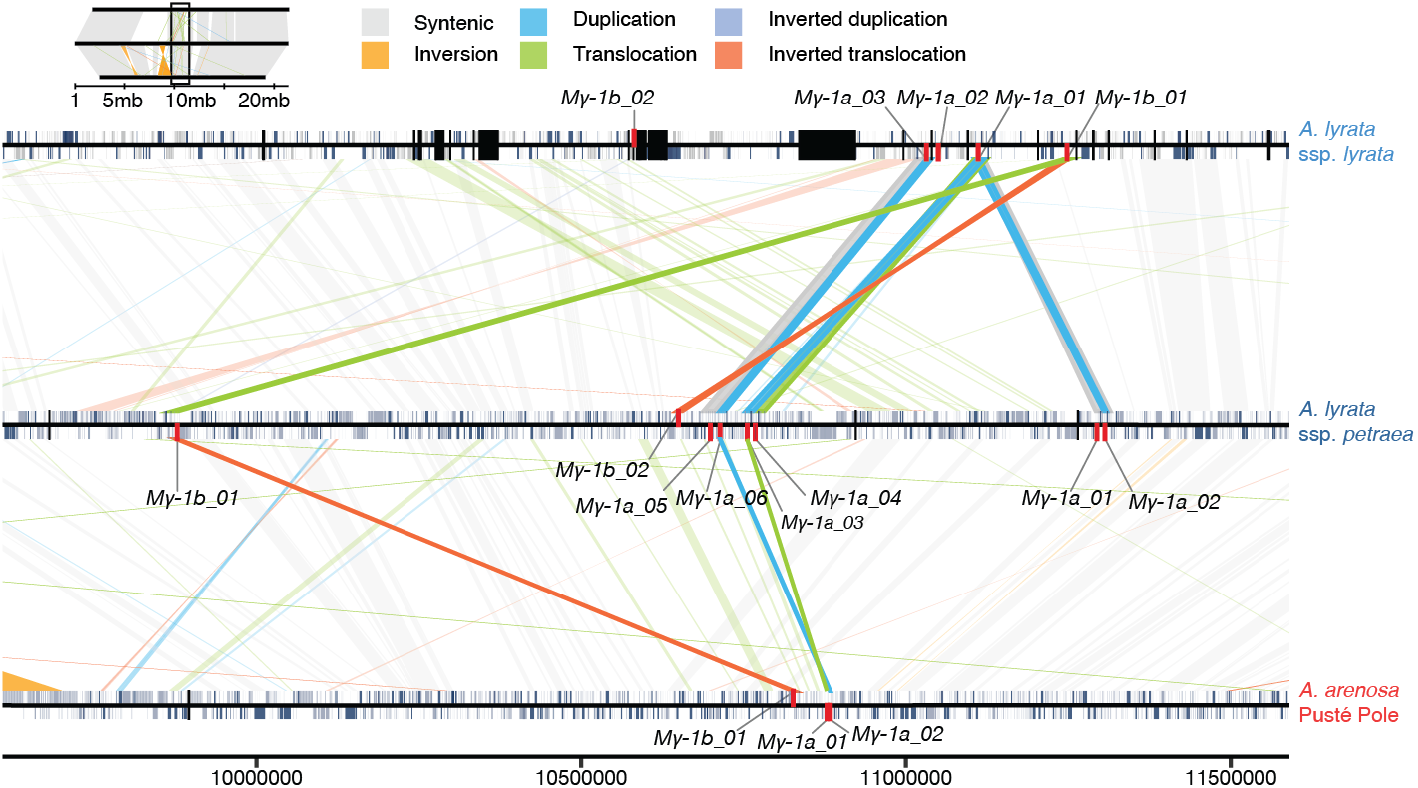
*Mγ-1* duplications in *Arabidopsis lyrata* ssp. *lyrata*, *A. lyrata* ssp. *petraea*, and *A. arenosa.* Localization and transpositions of *Mγ-1* genes (red) on segments of chromosome 2 in *A. lyrata* ssp*. lyrata*, *A. lyrata* ssp. *petraea* and *A. arenosa* Pusté Pole, with genes marked in blue, repeats in gray and assembly gaps in black. Syntenic sequences between assemblies are indicated by light gray connections. Inversions (orange), duplications (blue), translocations (green), inverted duplications (light purple), and inverted translocations (orange-red) were classified by SyRI. Scale bar and nucleotide positions refer to the *A. lyrata* ssp. *petraea* assembly.

To exclude artifacts, we compared the *A. lyrata* ssp. *petraea Mγ-1a* genomic loci with *A. lyrata* ssp. *lyrata*, and our primary and scaffolded assemblies supported this area well. Both genome assemblies aligned well through this region. We addressed this area’s syntenic relation, the origin, and the underlying nature of these gene duplications. *Mγ-1a* is represented by 3, 6, and 2 orthologs in *A. lyrata* ssp. *lyrata, A. lyrata* ssp. *petraea* and *A. arenosa* Pusté Pole, respectively (Figure 5).

We used SyRI synteny analysis to investigate the structural variants and the genomic context surrounding these orthologs close to the centromeric region of chromosome 2. The analysis show that the *A. lyrata* ssp. *lyrata Mγ-1a_*01 is quadrupled in *A. lyrata* ssp. *petraea,* creating *A. lyrata* ssp. *petraea Mγ-1a_01, Mγ-1a_02*, *Mγ-1a_03* and *Mγ-1a_04*. The *A. lyrata* ssp. *lyrata Mγ-1a_*03 is duplicated generating *Mγ-1a_05* and *Mγ-1a_06* in *A. lyrata* ssp. *petraea*. The *A. lyrata* ssp. *lyrata Mγ-1a_02* is structurally related to its paralog *Mγ-1a_01 in A. lyrata* ssp. *lyrata*, but exhibits no syntenic relation to *A. lyrata* ssp. *petraea*. The two *Mγ-1a* paralogs in *A. arenosa* Pusté Pole, *Mγ-1a_01* and *Mγ-1a_02* have a syntenic relation to *Mγ-1a_03* and *Mγ-1a_06* of *A. lyrata* ssp. *petraea,* respectively (Figure 5). Alignments of genomic sequences surrounding *Mγ-1* loci were generated using MAFFT, showing that the downstream repeat-rich area responsible for imprinting of *A. thaliana Mγ-1a (Makarevich et al., 2008)* can be found in one *A. arenosa* Pusté Pole (*Mγ-1a_01*) and four *A. lyrata* ssp. *petraea* loci (*Mγ-1a_01, Mγ-1a_02, Mγ-1a_04 and Mγ-1a_06*) (Figure S12). The presence of the repeat-rich area, and thereby duplication of the regulatory region, indicates that these genes may be imprinted in a similar manner as *PHE1 (Makarevich et al., 2008)*. However, imprinting studies in *A. lyrata* ssp. *petraea* is required to demonstrate this, since a repeat-rich region is also detected in one *A. lyrata* ssp. *lyrata* paralog, but no *Mγ-1a* imprinting was identified in a previous study searching for imprinting in *A. lyrata* ssp. *lyrata* (Klosinska *et al*., 2016).

To further analyze the functional regulation of the *Mγ-1* group, we used *A. arenosa* and *A. lyrata* ssp. *petraea* genome resources to identify the orthologous genes of previously described *Mγ-1a (PHE1)* transcription factor targets in *A. thaliana* (Batista *et al*., 2019). These *PHE1* targets were previously clustered into three groups in *A. thaliana* depending on their expression in the seed. Characteristic for cluster 1 is the down-regulation in *A. thaliana* during endosperm cellularization, whereas the two other clusters show only minor differential expression changes (Batista *et al*., 2019). Our differential expression analysis of *A. arenosa* and *A. lyrata* ssp. *petraea* orthologous *Mγ-1a (PHE1)* targets before and after endosperm cellularization demonstrated a strong down-regulation of cluster 1 *Mγ-1a* targets in *A. arenosa* Pusté Pole seeds (Figure 6A). In contrast, this drop of expression could not be detected in *A. lyrata* ssp. *petraea* cluster 1 *Mγ-1a* targets, where the expression level was relatively constant (Figure 6A). In summary, cluster 1 *Mγ-1a* targets are differentially regulated in *A. thaliana* and *A. arenosa* versus *A. lyrata* ssp. *petraea* during endosperm cellularization.

**Figure 6.**
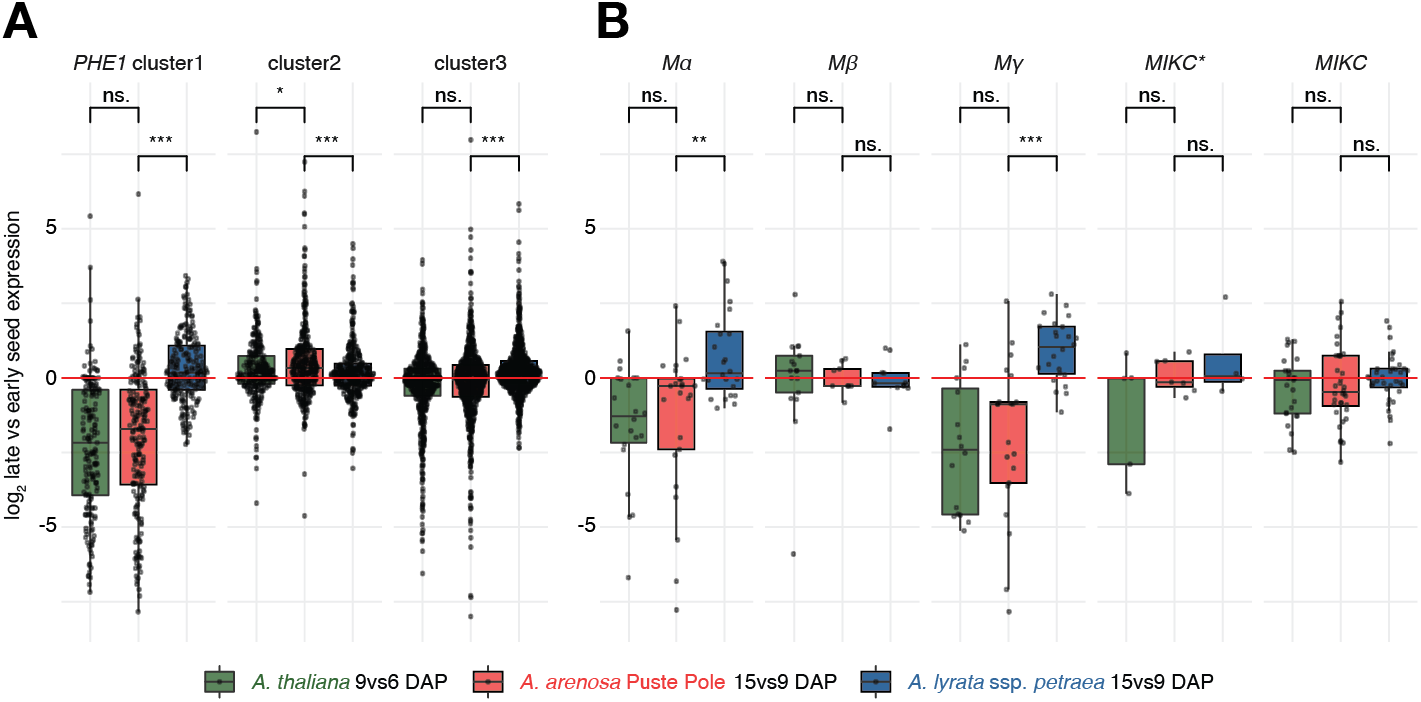
Differential expression of orthologs of *Mγ-1a PHE1*-targets and MADS-box genes before and after endosperm cellularization. (A) *Mγ-1a PHE1*-targets in *A. thaliana* were clustered depending on their expression during seed development (Batista *et al*., 2019). Characteristic for cluster 1 is the down-regulation in *A. thaliana* during endosperm cellularization (green). Differential expression of orthologs of *Mγ-1a PHE1*-targets in *A. arenosa* Pusté Pole and *A. lyrata* ssp. *petraea* are plotted in red and blue, respectively. Note that repression in cluster 1 is also seen in *A. arenosa* (red) but not in *A. lyrata* ssp. *petraea* (blue). (B) Differential gene expression of MADS-box genes before and after endosperm cellularization in *A. thaliana, A. lyrata* ssp. *petraea* and *A. arenosa* Pusté Pole. Similar to potential orthologs of *Mγ-1a PHE1*-targets, *Mα* and *Mγ* MADS-box type I gene expression declines around endosperm cellularization in *A. thaliana* (green) and *A. arenosa* (red) while increases in *A. lyrata* ssp. *petraea* (blue). DAP, days after pollination. Significance, ** p<0,01;*** p<0,001; ^n.s.^ not significant (Wilcoxon signed-rank test).

Focusing on the MADS-box type I genes, Mα and Mγ genes are down-regulated in *A. arenosa* Pusté Pole and *A. thaliana*, whereas in *A. lyrata* ssp. *petraea* seeds the orthologous Mα and Mγ genes are upregulated (Figure 6B).

The Mβ genes do not show significant differences. The substantial differences in Mα and Mγ expression between *A. arenosa* and *A. lyrata* ssp. *petraea* before and after cellularization indicate that these species differ in the regulation of the developmental timing of endosperm development. *A. thaliana* Mα and Mγ expression is regulated similarly to *A. arenosa.* In hybrid seeds between the species, endosperm developmental failure plays an important role in the establishment of species barriers (Lafon-Placette *et al*., 2017; Bjerkan *et al*., 2020), and the observed expression difference between the species may be part of the genetic mechanism leading to the species barrier in hybrid seeds.

## Conclusion

Here we have investigated and compared MADS-box type-I gene evolution in the genus *Arabidopsis* to gene expression in seed stages before and after endosperm cellularization. To do so we generated chromosome-scale assemblies of *A. lyrata* ssp. *petraea* and *A. arenosa*.

The quality of the reference genomes is essential for knowledge transfer from model species such as *A. thaliana* to their relatives, and research in the genus *Arabidopsis* has been supported by recent genome sequencing (Rawat *et al*., 2015; Hu *et al*., 2011) and improved annotation of the North American *A. lyrata* ssp. *lyrata*. This genome has served as reference for functional, ecological, and evolutionary experiments in *A. lyrata* and also *A. arenosa* (Klosinska *et al*., 2016; Yant *et al*., 2013; Arnold *et al*., 2016; Lafon-Placette *et al*., 2017; Bjerkan *et al*., 2020)*. Arabidopsis lyrata* has also been used as the reference for *Arabidopsis* reference-guided assemblies (Paape *et al*., 2018; Burns, Mandáková, Gunis, *et al*., 2021; Kolesnikova *et al*., 2023; Jaegle *et al*., 2023). The genus Arabidopsis has been extensively sequenced, e.g. *A. lyrata* ssp. *petraea (Akama et al., 2014; Paape et al., 2018)*, *A. halleri* (Briskine *et al*., 2016; Legrand *et al*., 2019), *A. kamchatica* (Paape *et al*., 2018), *A. suecica* (Novikova *et al*., 2017; Burns, Mandáková, Jagoda, *et al*., 2021; Jiang *et al*., 2021), and *A. arenosa* (Liu *et al*., 2020; Bohutínská *et al*., 2021; Burns, Mandáková, Jagoda, *et al*., 2021; Barragan *et al*., 2021). The assembly presented is the only fully *de novo* assembly of *A. arenosa* long reads, and only the fourth chromosome-level genome of any *Arabidopsis* species, to be published and released.

The genome assemblies allowed interspecies comparisons of MADS-box type-I orthologs, gene family expansion analysis and gene expression studies comparing orthologs. We demonstrate using MEME analysis identified motifs that are diagnostic for MADS-box type I clades, an additional guide for MADS-box type I classification but also allowing for future studies of the roles of these motifs. Our analysis suggests that chromosomal localisation may explain the high duplication rate of MADS-box type I clades in *A. lyrata* and *A. arenosa* compared to *A. thaliana*. Gene expansion is observed in orthologs located in centromeric regions whereas the same orthologs in a different species do not expand when located on chromosome arms. A negative correlation between gene duplication in MADS-box type I genes and seed-specific expression of these loci was observed, i.e. highly duplicated genes were associated with low expression values. Furthermore, the duplication rate was linked to previously identified MADS-box interaction (Parenicová *et al*., 2003; Qiu and Köhler, 2022), where highly duplicated genes were associated with less interactions. We also observed substantial differences in type-I Mα and Mγ expression between *A. arenosa* and *A. lyrata* ssp. *petraea* before and after endosperm cellularization indicating major differences in developmental regulation in the endosperm between the two species, suggesting a genetic cause for the endosperm based hybridization barrier developmental failure in *A. arenosa* and *A. lyrata* ssp. *petraea* hybrid seeds.

## Experimental procedures

### Plant lines and growth conditions

*Arabidopsis thaliana* accessions were obtained from the Nottingham Arabidopsis Stock Centre (NASC). The *A. arenosa* population MJ09-4 originates from Nízke Tatry Mts.; Pusté Pole (N 48.8855, E 20.2408) and *A. lyrata* ssp. *petraea* population MJ09-11 originates from lower Austria; street from Pernitz to Pottenstein (N 47.9190, E 15.9755) (Jørgensen *et al*., 2011; Lafon-Placette *et al*., 2017; Bjerkan *et al*., 2020). Seed sterilization was performed by washing in three steps with 70% ethanol, bleach solution (20% Klorin (Lilleborg Industrier), 0.1% Tween20), and wash solution (0.001% Tween20) or by over-night chlorine gas sterilization (Lindsey *et al*., 2017). Sterile seeds were planted on 0.5 Murashige and Skoog (MS) plates (Murashige and Skoog, 1962) supplemented with 2% sucrose. Seeds were stratified for 1 to 3 weeks at 4°C before transfer to growth chambers at 18°C long-day conditions (16 h light, 160 lmol/m2/sec, relative humidity 60–65%). Seedlings were transferred to soil for vegetative growth and vernalized at 9°C for 3 weeks to induce flowering.

### DNA isolation

DNA was isolated from one individual of *A. arenosa* Pusté Pole and one individual of *A. lyrata* ssp. *petraea*. All available flowers were harvested as input tissue, which resulted in approximately 1-1.5 g of tissue for each plant. Two different protocols were used: one protocol tailored for Illumina short reads, the other for PacBio Sequel I long reads. In both cases, the extraction protocols were the same for both species. For Illumina sequencing, DNA was isolated with the Ezna Plant DNA isolation kit (Omega Bio-tek). To extract high-molecular-weight (HMW) genomic DNA suitable for long-read sequencing, we first performed nuclei isolation (Rachael Workman *et al*., 2021), followed by the Nanobind Plant Nuclei Big DNA kit from Circulomics to isolate high molecular-weight DNA. For the isolation of nuclei the centrifuge speed used was 3800 g and the DNA isolation was performed using twice the amount of proteinase K and PL1 buffer, two nanodisks for DNA binding, and three washing steps using PW1. DNA purity and concentrations were checked with a NanoDrop ND-1000 and a Qubit 3 fluorometer (ThermoFisher). Fragment length was verified by running 2-5 µl of the isolate on a 0.5% agarose gel overnight at 30V. Four samples with 2 g of plant tissue including stem, flowers, and leaves were flash-frozen for Hi-C sequencing.

### Library preparation and sequencing

For both species, 5 μg of HMW DNA was used to generate a 20 kb library according to the manufacturer’s instructions (Pacific Biosciences, USA). The libraries were sequenced on PacBio Sequel I using Sequel Polymerase v3.0 and Sequencing Chemistry v3.0. Two SMRT-cells were sequenced for A. arenosa Pusté Pole and three SMRT-cells for A. lyrata ssp. petraea. Loading was performed by diffusion and the movie time was 600 min. Sequencing yielded 1,236,862 reads for *A. arenosa* (estimated coverage 86x), and 1,468,776 reads for *A. lyrata* (estimated coverage 72x). Sequencing on Illumina HiSeq 4000 resulted in 101,609,762 pair-end reads for *A. arenosa* (estimated coverage 169x) and 80,006,646 pair-end reads for *A. lyrata* (estimated coverage 120x). Crosslinked Hi-C DNA sequenced on Illumina HiSeq 4000 resulted in 308,885,555 paired reads (estimated coverage 515x) (Data S1.1). All sequencing was performed at the Norwegian Sequencing Center (NSC; https://www.sequencing.uio.no/).

### Genome size estimation

Mercury v1.0 (Rhie *et al*., 2020) was used to create a k-mer frequency spectrum of the Illumina reads with a k-mer length of 19. High-coverage (≥1000) k-mers were observed to be enriched in ribosomal, mitochondrial, and chloroplast DNA. GenomeScope 2.0 (Ranallo-Benavidez *et al*., 2020) was used to predict genome size and heterozygosity based on the k-mer spectra. Following the vertebrate genome project (Rhie *et al*., 2021), all k-mers were included. Relative fluorescence intensities were estimated by flow cytometry (FCM) using fresh plant tissue (Dolezel *et al*., 2007). *Solanum pseudocapsicum L.* (2C = 2.59 pg) was used as an internal standard. Relative fluorescence intensity of at least 3000 nuclei was recorded using a Partec Space flow cytometer (Partec GmbH, Münster, Germany) equipped with the UV-LED chip (365 nm). FCM results were expressed as fluorescence intensities relative to unit fluorescence intensity of the internal reference. The estimated haploid genome sizes are comparable to our flow cytometric measurements (Data S2) and estimates in previous studies (Pellicer and Leitch, 2020). The genome size estimates were used to estimate sequencing read coverage.

### De novo genome assembly

Six strategies were used to generate a variety of assemblies. One assembly per species was selected for further analysis. Long reads were repeatedly assembled with either Canu v2.1 (Koren *et al*., 2017), FALCON-unzip v1.3.7 (Chin *et al*., 2016), or Flye v2.6 (Kolmogorov *et al*., 2019). Long and short reads were assembled with MaSuRCA v3.3.5 (Zimin *et al*., 2013). Long reads were also polished with short reads using *LoRDEC* (Salmela and Rivals, 2014), then repeatedly assembled with Canu or Flye. All assemblies were computed on the Saga supercomputer (https://documentation.sigma2.no/hpc_machines/saga.html). Assemblies were compared by total length, N50-type statistics, longest contig, and gene content (BUSCO versions 3.0.2 and 4.14 (Manni *et al*., 2021; Simão *et al*., 2015; Waterhouse *et al*., 2018) using embryophyta_odb9 with 1440 genes from OrthoDB (https://www.orthodb.org/)). For both species, the Canu long-read contig assembly was selected for further processing.

### Polishing and haplotig phasing

The contig assemblies were polished using Illumina short reads from the same individual plants with 9 to 12 iterations of Pilon v1.23 (Walker *et al*., 2014). BUSCO (Manni *et al*., 2021; Simão *et al*., 2015; Waterhouse *et al*., 2018) found high inclusion rates but also high duplication rates for genes expected to be single copy (Figure S1 and Figure S3, indicating that the assembly might have separated both haplotypes of portions of the diploid genomes. Following the documentation on https://github.com/broadinstitute/pilon/wiki, the Purge Haplotigs v1.1.1 (Roach *et al*., 2018) and Purge_Dups v1.2.3 (Guan *et al*., 2020) pipelines were run to remove alternate haplotigs. For quality assessment, see Figure S1 to S5, and Data S1.

### Reference-guided scaffolding

The *A. lyrata* ssp. *petraea* assembly was scaffolded using RaGOO v1.11 (Alonge *et al*., 2019) guided by the published *A. lyrata* ssp. *lyrata* assembly (Hu *et al*., 2011). Manual curation of the computational results revealed the perpetuation of a chloroplast insertion on scaffold 2 and a misassembly on scaffold 1 of the reference (Slotte *et al*., 2013; Henry *et al*., 2014; Burns, Mandáková, Jagoda, *et al*., 2021). The computation had inserted a chloroplast-like contig in the first case, and broken several contigs in the second. The contigs and scaffolds were repaired manually in both cases.

### Hi-C scaffolding

Hi-C data for *A. arenosa* were generated with Arima kits and Illumina sequencing. The Hi-C reads were mapped to the draft assemblies with BWA-mem v0.7.17 (Li, 2013; Li and Durbin, 2009) and filtered with matlock (https://github.com/phasegenomics/matlock, commit 9fe3fdd). Scaffold candidates were generated by three algorithms: SALSA2 (Ghurye *et al*., 2019) https://github.com/marbl/SALSA (Ghurye et al., 2019)commit ed76685(Ghurye et al., 2019), FALCON-Phase vBeta Update 1 (Kronenberg *et al*., 2021), and ALLHiC (Zhang *et al*., 2019) https://github.com/tangerzhang/ALLHiC (Zhang *et al*., 2019)commit ffaa10e(Zhang *et al*., 2019) (Data S1). The FALCON-Phase scaffolding did not increase genome contiguity significantly and was excluded from further analysis (see Data S1). Scaffolded assemblies were compared to each other by constructing whole genome alignments between the newly scaffolded genomes, as well as to the *A. lyrata* ssp. *lyrata* reference genome (Figures S4 and S5). Scaffolds created by SALSA2 and ALLHiC were visualized in Juicebox (Durand *et al*., 2016) and curated with the Juicebox Assembly Tools (JBAT, https://github.com/aidenlab/Juicebox, commit 12bc674) (Figure S6) (Dudchenko *et al*., 2018). The JBAT version of the ALLHiC scaffolds was selected for further processing. The scaffolds representing both species were processed in the PBJelly gap-filling pipeline v15.8.24 (English *et al*., 2012).

### Repeat classification and masking

One of the largest influences on the quality of the gene prediction and the number of predicted genes was the initial RepeatModeler repeat masking step, which was carefully tuned to known repeat and genome models in *Arabidopsis*. The RepeatModeler v2.0.1 pipeline was run for *de novo* identification of transposable elements (TEs) (Flynn *et al*., 2020). This included TRF, RepeatScout, RECON TE detection, LTRharvest and LTR_retriever. Results were merged, clustered, and deduplicated. Repeats were classified by comparison to Dfam v3.1 (Storer *et al*., 2021). False positives were removed based on sequence homology (blastn ≤1e-10) to an *A. thaliana* cds database (Araport11_cds_20160703) (Cheng *et al*., 2017), which was beforehand cleaned for sequences with high sequence homology to known TEs in Brassicales (Dfam_3.1 database, (Storer *et al*., 2021)). The *de novo* predicted TE families were combined with the Dfam_3.1 and RepBase-20170127 databases and used in RepeatMasker v4.0.9 to annotate and soft mask the assemblies before gene annotation (Smit *et al*., 2015). Centromeric, ribosomal and telomeric repeats were identified and labeled using previously described sequences (Kawabe and Charlesworth, 2006; Maheshwari *et al*., 2017; Jin *et al*., 2020; Rhie *et al*., 2021).

### Genome annotation

The gene prediction was based on the BRAKER v2.1.5 pipeline, which trains GeneMark-EX and AUGUSTUS, including extrinsic evidence from RNA-seq and protein homology (Hoff *et al*., 2016; Hoff *et al*., 2019; Brůna *et al*., 2021). We used available RNA-seq for *A. arenosa* Pusté Pole and *A. lyrata* ssp. *petraea* from the Sequence Read Archive (Leinonen *et al*., 2011) and complemented this with seed transcriptomes (see below). In addition, the *A. thaliana* proteome was aligned by GenomeThreader (Gremme *et al*., 2005) to improve our species-specific training further and guide the gene prediction.

### Genome quality assessment

After each assembly processing step, quality control was used to assess our results and change or choose software and parameters. Primary genome statistics were determined by QUAST v5.0.2 (Gurevich *et al*., 2013). Feature Response Curve (FRC) was used to compare sequence quality (https://github.com/vezzi/FRC_align). Merqury v1.3 gave a k-mer based approach to sequence quality, sequence completes, and duplication rate (Rhie *et al*., 2020). BUSCO v4.14 (embryophyta_odb9) was used to monitor gene completeness and duplication rate (Manni *et al*., 2021). The structural variation was addressed using minimap2 (Li, 2018) alignments between assemblies and also the *A. lyrata* ssp. *lyrata* reference genome (Li, 2018). The alignments were further inspected with Assemblytics (Nattestad and Schatz, 2016) and dotPlotly (https://github.com/tpoorten/dotPlotly, commit 1174484) for visualization.

### Orthology prediction and synteny analysis

Orthogroups were inferred between *A. arenosa* Pusté Pole, *A. lyrata* ssp. *petraea*, and *A. thaliana* following the OrthoFinder v2.5.2 tutorials (Emms and Kelly, 2019). For synteny and collinearity assignment of homologous regions, MCScanX (Wang *et al*., 2012) was used with default parameters. Syntenic relations were further plotted with the RIdeogram package v0.2.2 (Hao *et al*., 2020). In addition, sequence differences were compared by whole-genome comparisons, and structural rearrangements were classified by SyRI vV1.5 (Goel *et al*., 2019).

### Identification of MADS-box genes

In addition to the two genomes produced in this study, we downloaded publicly available high-quality genome assemblies from the genus *Arabidopsis* from the National Center for Biotechnology Information (NCBI, https://www.ncbi.nlm.nih.gov/) and Phytozome 13 (https://phytozome-next.jgi.doe.gov/). The minimum requirement for the assemblies was a BUSCO completeness score above 95% and high contig continuity (i.e. near chromosome length assemblies). As an outgroup to *Arabidopsis,* we added genomes of *Capsella rubella* and *C. grandiflora* (Data S1) with a similar requirement for BUSCO-score and contig continuity. For genome assemblies without previously predicted genes, we used AUGUSTUS with the *-arabidopsis* gene model.

We then constructed a database consisting of the known AGLs (AGAMOUS-like genes) from *A. thaliana* (108 in total) retrieved from the Arabidopsis Information Resource (TAIR, https://www.arabidopsis.org/). AGL26 (At5g26880) was excluded from the database since this is a gene coding for an RNA methyltransferase and does not have a MADS-box. In addition, we identified five MADS-box genes from the PlantTFDB website (http://planttfdb.cbi.pku.edu.cn/) which do not currently have an AGL designation in *A. thaliana*. For the list of the genes with their corresponding AGL names as well as commonly used names and abbreviations in *A. thaliana*, see Data 4.1. Previously identified MADS-box genes from *A. thaliana* were added as a basic local alignment search tool (BLAST) database in Geneious 2020.2.4 (https://www.geneious.com).

The database of MADS box genes from *A. thaliana* was used in blastp searches (BLAST+ v2.12.0 (Altschul et al., 1990)) against the predicted genes of *A. lyrata* ssp. *petraea*, *A. arenosa* Pusté Pole, and three published *Arabidopsis* genomes that met our criteria: *A. lyrata* ssp. *lyrata* (https://phytozome-next.jgi.doe.gov/info/Alyrata_v2_1), *A. arenosa* Strecno (https://www.ncbi.nlm.nih.gov/assembly/GCA_902996965.1), and *A. halleri halleri* (https://www.ncbi.nlm.nih.gov/assembly/GCA_900078215.1), (Data S1.5). In addition, we blasted the predicted genes of two species of *Capsella* against the same database (*Capsella grandiflora*, https://phytozome-next.jgi.doe.gov/info/Cgrandiflora_v1_1 and *Capsella rubella,* https://www.ncbi.nlm.nih.gov/assembly/GCF_000375325.1/). The top five hits for each known AGL were kept for each of the species, with any duplicates removed. All sequences were then annotated with InterProscan (http://www.ebi.ac.uk/interpro implemented in Geneious Prime 2021) to verify the presence of the MADS-box region, and sequences with no MADS-box domain were removed. The results from the InterProscan v5.47 analysis were also used to identify potential mistakes in the predicted gene sequences which were manually corrected (Data S3.2).

### Phylogenetic analysis

A multiple sequence alignment was constructed from the blast results together with the canonical A*. thaliana* MADS-box genes from TAIR (https://www.arabidopsis.org) and PlantTFDB (http://planttfdb.cbi.pku.edu.cn/) with MUSCLE version 3.8.425 (Edgar, 2004). A phylogenetic tree was constructed based on the alignment with FastTree2 v2.1.11 (Price *et al*., 2010) and visualized in Geneious Prime 2021. The resulting tree was manually inspected for any outliers and long branches. Long-branched clades and sequences without any clear homologs to known MADS-box genes were pruned from the tree and removed from the alignment, the remaining sequences realigned, and a new tree was constructed with FastTree2. We repeated the process of tree building, pruning, and realignment until no more spurious clades remained in the tree. The resulting alignment with 841 amino acid sequences was refined with MUSCLE, trimmed with trimAl, and a final phylogenetic tree was constructed with IQ-tree v1.6.12 with ultrafast bootstrap approximation and SH-like approximate likelihood ratio test (Minh *et al*., 2020). The trees in Figure 2 and Supplemental Figure S8 were visualized in FigTree 1.4.4 (Rambaut 2016).

### Motif analysis with MEME

To identify conserved motifs between the MADS-box genes of the *Arabidopsis* genus, MEME version 5.1.1 (Multiple Expectation Maximization for Motif Elicitation, http://meme-suite.org/) was used. We scanned for the 50 most common motifs with lengths above 20 AA; a graphical representation for the relative position of the motifs on the sequences can be found in Figure S8. The full results, motif sequences and a html version can be found on https://github.com/PaulGrini/Arabidopsis_assemblies. The full-length sequences of the proteins were grouped based on the phylogenetic analysis and commonly occurring MEME motifs into Mα, Mβ, Mγ, MIKC*, and MIKC. A principal component analysis was performed on the occurrence of MEME motifs using the PCAtools package in R (Blighe K, 2022).

### Analysis of syntenic regions surrounding Mγ-1

Genomic sequences of *Mγ-1* and their surrounding upstream and downstream regions from *A. thaliana, A. lyrata* ssp. *petraea* and *A. arenosa* Pusté Pole were aligned with MAFFT v7.470 (Katoh and Standley, 2013) to identify syntenic regions and localize transpositions. SyRI (Synteny and Rearrangement Identifier, https://github.com/swbak/SyRI commit 29a9272 (Goel *et al*., 2019)) was used to detect and classify structural differences between genomes, i.e inversions, duplications, translocations, inverted duplications, and inverted translocations.

### Expression analysis of MADS-box genes in *A. arenosa* and *A. lyrata* compared to *A. thaliana*

Seeds of *A. lyrata* ssp. *petraea* and *A. arenosa* Pusté Pole were dissected from siliques at 9 and 15 DAP and shock frozen in liquid nitrogen. Around 20 seeds were pooled into one tube, and RNA was extracted from three to four replicates per plant and stage using the Spectrum Plant Total RNA kit (SIGMA). MagNA Lyser Green Beads (Roche) was used for the initial lysis step as described in Shirzadi et al. (Shirzadi *et al*., 2011). RNA concentration and quality were measured with a NanoDrop ND-1000, Bioanalyzer 2100 Agilent RNA 6000 Nano kit, and a Qubit 3 fluorometer (ThermoFisher) using the Qubit RNA BR Assay kit (Invitrogen). All kits were used according to the manufacturers’ instructions. Whole cDNA libraries were prepared by the NSC from total RNA using the Illumina TruSeq Standard mRNA Library Prep kit. The samples were sequenced on an Illumina (HiSeq 4000) sequencer creating 150 bp pair-end reads with 350 bp inserts. TrimGalore v0.4.4, a wrapper script around Cutadapt and FastQC (Krueger, 2016), was used for adapter and quality trimming. The reads were mapped using HISAT2 v2.2.1 (Kim *et al*., 2019) and default parameters and quantified by featureCounts (Liao *et al*., 2014) and DESeq2 library normalization (Love *et al*., 2014). MADS-box type I genes were selected, grouped, and ordered following the MADS-box phylogeny (Figure 2). The log2-transformed reads were displayed in a heat map drawn with the iheatmapr v0.5.2.9000 package (in Figure 4 and Figure 11) (N Schep and K Kummerfeld, 2017) including as a reference previously published *A. thaliana* seed expression samples (Bjerkan *et al*., 2020). The reads were reanalyzed using the same (described above) processing steps.

## Data availability

All sequences generated in this study have been deposited in the National Center for Biotechnology Information Sequence Read Archive (https://www.ncbi.nlm.nih.gov/sra) with project number PRJNA844220. The *A. lyrata* ssp. *petraea* genome is depostited at NCBI with the reference number GCA_026151145.1 (https://www.ncbi.nlm.nih.gov/datasets/genome/GCA_026151145.1/), the *A. arenosa* Pusté Pole genome has reference number GCA_026151155.1 (https://www.ncbi.nlm.nih.gov/datasets/genome/GCA_026151155.1/). An accompanying GitHub repository can be found at https://github.com/PaulGrini/Arabidopsis_assemblies. The repository contains predicted genes, scripts, and additional information for the analyses in this paper.

## Supporting information

Supplementary Data S1: Sequencing and assemblies

Supplementary Data S2: Genome size estimation

Supplementary Data S3: Sequence names for MADS-genes

Supplementary Figure S1: Pipeline for genome assembly

Supplementary Figure S2: GenomeScope profiles

Supplementary Figure S3: Copy number spectrum plots

Supplementary Figure S4: A. lyrata ssp. petraea - A. lyrata ssp. lyrata alignments

Supplementary Figure S7: Genome alignments

Supplementary Figure S8: Extended MADS-box phylogeny

Supplementary Figure S9: MADS-box type II phylogeny

Supplementary Figure S10: PCA of MEME motifs

Supplementary Figure S11: MADS-box type II expression during seed development

Supplementary Figure S12: MAFFT alignments of genomic sequences

## Acknowledgements

The sequencing service was provided by the Norwegian Sequencing Centre (www.sequencing.uio.no), a national technology platform hosted by the University of Oslo and supported by the “Functional Genomics” and “Infrastructure” programs of the Research Council of Norway and the Southeastern Regional Health Authorities. We thank the Laboratory of Flow Cytometry, Institute of Botany, Academy of Sciences (Czech Republic) for assistance.

## Short Legends for Supporting Information

**Figure S1:** Pipeline for genome assembly

**Figure S2:** GenomeScope profiles

**Figure S3:** Copy number spectrum plots

**Figure S4:** *A. lyrata* ssp*. petraea-*ssp. *lyrata* alignments

**Figure S5:** *A. arenosa* Pusté Pole*-A. lyrata* ssp. *lyrata* alignments

**Figure S6:** *A. arenosa* Pusté Pole Hi-C contact maps

**Figure S7:** Genome alignments indicate genomic rearrangements

**Figure S8:** Extended MADS-box phylogeny

**Figure S9:** Phylogenetic analysis of MADS-box type II genes in *Arabidopsis*

**Figure S10:** PCA of the distribution of MEME motifs on MADS-box genes

**Figure S11:** MADS-box type II expression during seed development

**Figure S12:** MAFFT alignments of genomic sequences containing and surrounding *Mγ-1*

**Data S1:** An overview of the sequencing effort for *A. arenosa* and *A. lyrata petraea*

**Data S2:** Flow cytrometry and k-mer estimation of genome size

**Data S3:** Sequence names for all MADS-genes included in the phylogeny

## Supplemental Figures

**Figure S1: Pipeline for genome assembly.** *A. lyrata* ssp. *petraea* (blue) and *A. arenosa* Pusté Pole (red). *A. arenosa* was assembled *de novo* from Pacbio long-read data and scaffolded using Hi-C data. For *A. lyrata* ssp. *petraea a draft assembly was made de novo from PacBio long-reads, which were subsequently* scaffolded against the published genome of *A. lyrata ssp. lyrata (Hu et al., 2011)*. (A) Pipeline indicating each step in the assembly process. For each step additional programs or methods were tested; details of these analyses can be found in Data S1. (B) Statistics used in quality control and software decisions. Sequence contiguity is shown in NG50, using the calculated genome size of 201,144,702 bp for *A. lyrata* ssp. *petraea,* and 179,232,250 bp for *A. arenosa* Pusté Pole. (C) BUSCO scores using the gene set for Embryophyta odb9 with 1440 single-copy genes. For the assembly steps, completeness and duplication can be seen in light and dark blue, respectively. The combination of PacBio long reads with Canu created near-complete assemblies. Allelic duplications were identified and removed using Purge_Dups. (D) Phred Quality Score (QV) was calculated with Merqury (Rhie *et al*., 2020). A QV of 30 corresponds to 99.9% accuracy and QV 40 to 99.99%. The quality of both long read assemblies was improved by the Pilon polishing step using Illumina reads. While the *A. lyrata* ssp. *petraea* assembly shows a consistently lower error rate, both assemblies have high quality during all assembly steps. For more assemblies and detailed statistics see Data S1.

**Figure S2: GenomeScope profiles.** Mercury produced the k-mer frequency spectra of Illumina reads (blue) for a k-mer length of 19 nt. To this spectrum, the GenomeScope model (black line) was fitted, and genome length, repetitiveness, and heterozygosity rates were predicted. (A) K-mer spectrum for *A. lyrata* ssp. *petraea*. GenomeScope v2.0 infers a genome length of 201,144,702 bp with a heterozygosity rate of 1.46%. The heterozygous and homozygous peaks were detected with an approximal coverage of 50 and 100, respectively. K-mers with coverage above 1000 were enriched for ribosomal, mitochondrial, and chloroplast DNA. (B) K-mer spectrum for *A. arenosa* Pusté Pole. The heterozygous and homozygous peaks were detected with an approximal coverage of 50 and 100, respectively. GenomeScope infers a genome length of 179,232,250 bp with a heterozygosity rate of 1.61%.

**Figure S3: Copy number spectrum plots.** For quality control, the k-mer frequencies from every assembly were compared to the k-mer frequencies of the corresponding Illumina reads. The k-mer spectrum shows how many unique 19k-mer exist (Count) with a specific coverage (k-mer multiplicity). Each Illumina read k-mer is further grouped and colored depending on how often it is found in the assemblies. The first peak at half coverage is expected to contain k-mers found only on one haplotype, while the second peak should include k-mer from both haplotypes. (A) Copy number spectrum plot (spectra-cn) for the primary *A. lyrata* ssp. *petraea* assembly. The assembly was created with the Canu assembler using PacBio long reads. The spectrum indicates a high-quality assembly where nearly all k-mers found in the Illumina reads (not used for the assembly) were also detected in the assembly with their expected frequencies. K-mers found only in the long-read assembly are displayed as a red/blue-bar to the left of the spectrum plot. (B) Spectra-cn plot for the primary *A. arenosa* Pusté Pole assembly. The Canu assembler using uncorrected PacBio long-reads created the most complete draft assembly. Slightly reduced haploid resolution compared to the *A. lyrata* ssp. *petraea* assembly. (C) Spectra-cn plot of the *A. lyrata* ssp. *petraea* assembly after the Pilon polishing step. Using Illumina reads for polishing reduced the number of suspected erroneous k-mers found only in the assembly (blue/red bar on the left). (D) Spectra-cn plot of the *A. arenosa* Pusté Pole assembly after the Pilon polishing step. (E) Spectra-cn plot of the *A. lyrata* ssp. *petraea* assembly after the haplotic purging step. Only the primary haplotig is compared to the Illumina read set. (F) Spectra-cn plot of the *A. arenosa* Pusté Pole assembly after the haplotig purging. K-mer frequencies for the primary haplotig are shown. Half of the single-copy k-mers are missing and are found in the alternative haplotig contigs.

**Figure S4: *A. lyrata* ssp*. petraea-*ssp. *lyrata* alignments.** After each processing step, all *A. lyrata* ssp*. petraea* contigs and scaffolds were aligned to the *A. lyrata* ssp. *lyrata* reference genome as quality control. Larger mis-assemblies were easily detected and excluded. The reference scaffolds are on the x-axis, while the respective new assemblies are sorted on the y-axis. (A) Alignment of the *A. lyrata* ssp. *petraea* Canu draft assembly. (B) Alignment of the contigs after the Pilon polishing step shows a high duplication rate compared to the haploid reference. (C) Alignment of the contigs after the haplotic purging step. Contiguity remained after the removal of duplicated sequences. (D) Reference alignment after the reference-based scaffolding step using RaGoo. (E) Alignment after the gab closure by PBJelly displays no considerable changes. (F) Alignment of our final curated *A. lyrata* ssp*. petraea* assembly. The rearrangement of the previously reported misassembly for *A. lyrata* ssp. *lyrata* can be seen between scaffolds one and two (sorted by size) in the top left corner.

**Figure S5: *A. arenosa* Pusté Pole*-A. lyrata* ssp. *lyrata* alignments** For quality control, all *A. arenosa* Pusté Pole sequences were aligned to the *A. lyrata* ssp. *lyrata* reference genome. Although structural variation is expected between the species, inconsistent rearrangements between assemblies were fast detected and excluded. The y-axis represents the length sorted scaffolds from the published *A. lyrata* ssp. *lyrata* genome. (A) Contigs of the *A. arenosa* Pusté Pole Canu draft assembly on the y-axis show a high number of overlaps. (B) Alignment of the same contigs after the Pilon polishing step. (C) Alignment of the contigs after the haplotic purging step. Only the primary haplotig is shown. Contiguity remains after the removal of duplicated sequences. (D) The alignment plots were especially helpful to remove erroneous Hi-C scaffolding approaches. The best performance displayed here is our selected approach using ALLHiC. (E) Minor scaffolding mistakes were curated with the Juicebox Assembly Tools (JBAT) based on the Hi-C linkage map. The curated result aligned astoundingly well to the *A. lyrata* ssp. *lyrata* scaffolds. (F) Gap-closure with PBJelly did not change the scaffold arrangement. The previously reported misassembly on *A. lyrata* ssp. *lyrata* scaffold can be seen in the top left corner. In addition, one more extensive inversion is displayed on scaffold 7, in the center of the plot.

**Figure S6: *A. arenosa* Pusté Pole Hi-C contact maps.** Hi-C contact heat maps indicate the number of contacts between any given pair of loci in the assembly (red scale). Scaffolds are indicated by a blue line, and the green line documents the manual separation used for rearrangements. (A) Hi-C assembly heat map for *A. arenosa* Pusté Pole produced by the SALSA scaffolder. Strong signals far away from the diagonal were used for further scaffolding improvements. (B) The resulting contact heat map of the JBAT curated SALSA scaffolds. (C) Hi-C map of the *A. arenosa* Pusté Pole assembly scaffolded by the ALLHiC software. Problematic sequences are indicated by the low number of connections and stronger far-away signals. (D) Contact matrix after JBAT curation of miss joints and inversions. Green triangles indicate scaffold brakes. This assembly was selected and finalized by gap-filling with PBJelly (Figures S5 E and F).

**Figure S7: Genome alignments indicate genomic rearrangements.** SyRI (Synteny and Rearrangement Identifier) was used to detect and classify structural differences between genomes. (A) Comparison of our *A. lyrata* ssp. *petraea* assembly (dark blue) to the *A. lyrata* ssp. *lyrata* reference (light blue). Syntenic regions are shown as gray blocks, while structural differences are grouped in translocations (green), inversions (orange), and duplications (blue). For visualization purposes, the relocations between non-homolog scaffolds have been excluded (e.g., for translocation between scaffolds one and two, see Figure 1E). (B) Synteny alignment between *A. lyrata* ssp*. petraea* (dark blue) and *A. arenosa* Pusté Pole (red). Structural differences are indicated between largely syntenic scaffolds.

**Figure S8: Extended MADS-box phylogeny.** Phylogeny of all 841 identified MADS-box genes in *Arabidopsis* and *Capsella*. The groups correspond to previously published results with type I genes divided into three main groups (Mα, Mβ, Mγ), while type II genes fell into two monophyletic groups, MIKC and MIKC* [also referred to as Mδ] (Henschel *et al*., 2002; Parenicová *et al*., 2003; Arora *et al*., 2007; Thangavel and Nayar, 2018; Gramzow and Theißen, 2013; Qiu and Köhler, 2022). The groups are colored with a red box around Mα, blue around Mβ green around Mγ, yellow around MIKC*, and grey around MIKC. Sequences of *A. thaliana* are marked in blue, and the outgroup *Capsella* in orange. Monophyletic groups of *Arabidopsis* MADS-box genes, sharing a common ancestor with genes from the outgroup *Capsella,* are numbered starting from the top of the tree and delineated with solid lines. Additional gene duplications that occurred in the common ancestor of *Arabidopsis*, but after *Arabidopsis* and *Capsella* separated, are delineated with a dashed line and marked with an additional letter. For instance, “Mγ-1a” and “Mγ-1b” indicate that these two clades share a last common ancestor with *C. rubella* (i.e. “Mγ-1”), but have a gene duplication specific for *Arabidopsis* that occurred after the separation of *Arabidopsis* and *Capsella* . The right-hand column contains the result of the MEME analysis for each sequence. For the full result of the MEME analysis consult the material on GitHub.

**Figure S9: Phylogenetic analysis of MADS-box type II genes in *Arabidopsis*.** The tree was derived by a maximum likelihood analysis of 275 identified MADS-box type I sequences from *A. thaliana, A. lyrata* ssp. *petraea, A. arenosa* Strecno*, A. arenosa* Pusté Pole, and *A. halleri* with two species of *Capsella* (89 sequences; not shown in the figure) used as outgroup. Solid branches represent bootstrap support > 85%, while branches with support values < 85% are dashed. The root of the tree is placed between type I and type II genes (the corresponding tree and heatmap for type I genes can be found in Figure 2). Triangles represent clades where branches are collapsed at the most recent gene duplication event in the last common ancestor of the genus *Arabidopsis*. The length of the triangles corresponds to the overall branch length of the collapsed clade, see the main text for naming schemes of clades. The heatmap shows the number of gene copies for each clade in the genomes of *A. thaliana, A. lyrata* ssp. *petraea, A. arenosa* Strecno*, A. arenosa* Pusté Pole, and *A. halleri.* The column next to the heatmap indicates the canonical AGL names of the genes in *A. thaliana* found in the corresponding clade; “none” means that a gene representing the clade is not found in the *A. thaliana* genome, while “new” indicates that the gene does not have a given AGL name. The last column shows a simplified representation of the MEME motifs. A fully expanded phylogenetic tree with individual tip labels, support values for all branches, and the outgroup *Capsella* can be found in Figure S8. Results from the full MEME analysis can be found in Figure S8.

**Figure S10: PCA of the distribution of MEME motifs on MADS-box type I and type II genes.** The PCA was constructed from motifs identified by MEME and counted for each clade in the phylogeny. The first principal component axis captures 24.8% of the variation, and the second axis 10.2%. A polygon is drawn around each of the main groups with Mα in red, Mβ in blue, Mγ in green, MIKC* in yellow, and MIKC in grey.

**Figure S11: MADS-box type II expression during seed development.** Gene expression profiles are displayed for all identified MADS-box type II genes and compared between *A. thaliana* (right column), *A. arenosa* Pusté Pole (middle column), and *A. lyrata* ssp. *petraea* (left column). The *A. thaliana* development time series (Bjerkan *et al*., 2020) serves as a reference. The endosperm cellularization in *A. thaliana* occurs between the 6 and 9 days after pollination (DAP). To adjust for the relatively slower development in *A. arenosa* Pusté Pole and *A. lyrata* ssp. *petraea*, corresponding stages before and after endosperm cellularization were sampled at 9 and 15 DAP, respectively. Ortholog genes are grouped and ordered and follow our MADS-box phylogeny (Figure 2). Sample normalized counts are shown with a base-2 logarithmic scale. The MADS-box type II seed expressions remain rather constant compared to the strong decline of type I expression around endosperm cellularization (Figure 4). However, expression differences can be seen between the *Arabidopsis* species. The *SEPALLATA* (*MIKC-1/2/3*) genes are strongly expressed in *A. lyrata* ssp. *petraea* and *A. arenosa* seeds while only weakly or absent in *A. thaliana*. In addition to their largely functional redundant role in flower development and ovary formation (Pelaz *et al*., 2001; Kaufmann *et al*., 2009), a crucial role in fruit development and ripening has been reported for *SEPALLATA* orthologs in tomato, strawberry, and apple (Ampomah-Dwamena *et al*., 2002; Seymour *et al*., 2011; Schaffer *et al*., 2013). Furthermore, the orthologs of *MIKC-15* (*AGL19*), *MIKC-24* (*AGL16*), *MIKC-26* (*AGL24*) as well as the FLC clade *MIKC-28* to *MIKC-32* show strong expression differences reflecting the different regulations of flowering and strengthen of vernalization between the perennial *A. lyrata* ssp. *petraea* and *A. arenosa* plants compared to the annual *A. thaliana* (Schönrock *et al*., 2006; Alexandre and Hennig, 2008; Hu *et al*., 2014; Müller-Xing *et al*., 2022; Kemi *et al*., 2013; Soppe *et al*., 2021). We detect differing expressions of *MIKC*s* between the species. *MIKC*-2* and *MIKC*-3* (*AGL30* and *AGL65*) are expressed in *A. lyrata* ssp. *petraea* and *A. arenosa* seeds but are not detected in *A. thaliana*. On the other hand, *MIKC*-6* and *MIKC*-7* (*AT4G37435* and *AGL33*) show expression in *A. arenosa* and *A. thaliana* but not in *A. lyrata* ssp. *petraea* seeds. *MIKC*s* are known for their transcription activity in pollen (Verelst *et al*., 2007) but have also been detected in endosperm (Zhang *et al*., 2018). In addition, a high level of redundant heterodimers has been reported and double mutants reduce pollen fertility (Adamczyk and Fernandez, 2009).

**Figure S12: MAFFT alignments of genomic sequences containing and surrounding *Mγ-1*.** Protein coding sequences (CDS), which are displayed as dark blue arrows of *PHE1* orthologs from *A. thaliana* (green), *A. arenosa* (red), *A. lyrata* ssp. *lyrata* (light blue) and *A. lyrata* ssp. *petraea* (dark blue), are placed in the center. *A. thaliana* is highlighted by a light green background. On top, the consensus with identity score indicates a high similarity of the 3 prime regions downstream of the *Mγ-1* loci. Around 2200 bp 3 prime of *PHE1* lies the repeat-rich region crucial for its parental-specific expression (inside the AT1TE79790 RC/Helitron, highlighted by a transparent gray box). Similar locations of Line 1 (light green arrows) and LTR transposons (light blue arrows) can be found next to the *Mγ-1* loci between species. Further repeats are marked as light gray arrows and lncRNA as violet arrows (XR_002328948.1 and XR_002334149.1).

## References

1. Adamczyk, B.J. and Fernandez, D.E. (2009) MIKC* MADS domain heterodimers are required for pollen maturation and tube growth in Arabidopsis. Plant Physiol., 149, 1713–1723.

2. Akama, S., Shimizu-Inatsugi, R., Shimizu, K.K. and Sese, J. (2014) Genome-wide quantification of homeolog expression ratio revealed nonstochastic gene regulation in synthetic allopolyploid Arabidopsis. Nucleic Acids Res., 42, e46–e46.

3. Alexandre, C.M. and Hennig, L. (2008) FLC or not FLC: the other side of vernalization. J. Exp. Bot., 59, 1127–1135.

4. Alonge, M., Soyk, S., Ramakrishnan, S., Wang, X., Goodwin, S., Sedlazeck, F.J., Lippman, Z.B. and Schatz, M.C. (2019) RaGOO: fast and accurate reference-guided scaffolding of draft genomes. Genome Biol., 20, 224.

5. Ampomah-Dwamena, C., Morris, B.A., Sutherland, P., Veit, B. and Yao, J.-L. (2002) Down-regulation of TM29, a tomato SEPALLATA homolog, causes parthenocarpic fruit development and floral reversion. Plant Physiol., 130, 605–617.

6. Arnold, B.J., Lahner, B., DaCosta, J.M., Weisman, C.M., Hollister, J.D., Salt, D.E., Bomblies, K. and Yant, L. (2016) Borrowed alleles and convergence in serpentine adaptation. Proc. Natl. Acad. Sci. U. S. A., 113, 8320–8325.

7. Arora, R., Agarwal, P., Ray, S., Singh, A.K., Singh, V.P., Tyagi, A.K. and Kapoor, S. (2007) MADS-box gene family in rice: genome-wide identification, organization and expression profiling during reproductive development and stress. BMC Genomics, 8, 242.

8. Barragan, A.C., Collenberg, M., Schwab, R., Kerstens, M., Bezrukov, I., Bemm, F., Požárová, D., Kolář, F. and Weigel, D. (2021) Homozygosity at its Limit: Inbreeding Depression in Wild Arabidopsis arenosa Populations. bioRxiv, 2021.01.24.427284.

9. Batista, R.A., Moreno-Romero, J., Qiu, Y., Boven, J. van, Santos-González, J., Figueiredo, D.D. and Köhler, C. (2019) The MADS-box transcription factor PHERES1 controls imprinting in the endosperm by binding to domesticated transposons. Elife, 8, e50541.

10. Bemer, M., Heijmans, K., Airoldi, C., Davies, B. and Angenent, G.C. (2010) An atlas of type I MADS box gene expression during female gametophyte and seed development in Arabidopsis. Plant Physiol., 154, 287–300.

11. Bemer, M., Wolters-Arts, M., Grossniklaus, U. and Angenent, G.C. (2008) The MADS domain protein DIANA acts together with AGAMOUS-LIKE80 to specify the central cell in Arabidopsis ovules. Plant Cell, 20, 2088–2101.

12. Bjerkan, K.N., Hornslien, K.S., Johannessen, I.M., et al. (2020) Genetic variation and temperature affects hybrid barriers during interspecific hybridization. Plant J., 101, 122–140.

13. Blighe K, L.A. (2022) PCAtools: Everything Principal Components Analysis. R package version 2.10.0. *Github*. Available at: https://github.com/kevinblighe/PCAtools.

14. Bohutínská, M., Handrick, V., Yant, L., Schmickl, R., Kolář, F., Bomblies, K. and Paajanen, P. (2021) De-novo mutation and rapid protein (co-)evolution during meiotic adaptation in Arabidopsis arenosa. Mol. Biol. Evol., msab001–.

15. Briskine, R.V., Paape, T., Shimizu-Inatsugi, R., Nishiyama, T., Akama, S., Sese, J. and Shimizu, K.K. (2016) Genome assembly and annotation of Arabidopsis halleri, a model for heavy metal hyperaccumulation and evolutionary ecology. Mol. Ecol. Resour., 17, 1025–1036.

16. Brůna, T., Hoff, K.J., Lomsadze, A., Stanke, M. and Borodovsky, M. (2021) BRAKER2: automatic eukaryotic genome annotation with GeneMark-EP+ and AUGUSTUS supported by a protein database. NAR Genomics and Bioinformatics, 3, lqaa108–.

17. Burkart-Waco, D., Josefsson, C., Dilkes, B., Kozloff, N., Torjek, O., Meyer, R., Altmann, T. and Comai, L. (2012) Hybrid incompatibility in Arabidopsis is determined by a multiple-locus genetic network. Plant Physiol., 158, 801–812.

18. Burkart-Waco, D., Ngo, K., Dilkes, B., Josefsson, C. and Comai, L. (2013) Early disruption of maternal-zygotic interaction and activation of defense-like responses in Arabidopsis interspecific crosses. Plant Cell, 25, 2037–2055.

19. Burkart-Waco, D., Ngo, K., Lieberman, M. and Comai, L. (2015) Perturbation of parentally biased gene expression during interspecific hybridization. PLoS One, 10, e0117293.

20. Burns, R., Mandáková, T., Gunis, J., Soto-Jiménez, L.M., Liu, C., Lysak, M.A., Novikova, P.Y. and Nordborg, M. (2021) Gradual evolution of allopolyploidy in Arabidopsis suecica. Nat Ecol Evol, 5, 1367–1381.

21. Burns, R., Mandáková, T., Jagoda, J., Soto-Jiménez, L.M., Liu, C., Lysak, M.A., Novikova, P.Y. and Nordborg, M. (2021) Gradual evolution of allopolyploidy in Arabidopsis suecica. Nature Ecology & Evolution, 2020.08.24.264432.

22. Cheng, C.-Y., Krishnakumar, V., Chan, A.P., Thibaud-Nissen, F., Schobel, S. and Town, C.D. (2017) Araport11: a complete reannotation of the Arabidopsis thaliana reference genome. Plant J., 89, 789–804.

23. Chen, Z.J., Comai, L. and Pikaard, C.S. (1998) Gene dosage and stochastic effects determine the severity and direction of uniparental ribosomal RNA gene silencing (nucleolar dominance) in Arabidopsis allopolyploids. Proc. Natl. Acad. Sci. U. S. A., 95, 14891–14896.

24. Chin, C.-S., Peluso, P., Sedlazeck, F.J., et al. (2016) Phased diploid genome assembly with single-molecule real-time sequencing. Nat. Methods, 13, 1050–1054.

25. Colombo, M., Masiero, S., Vanzulli, S., Lardelli, P., Kater, M.M. and Colombo, L. (2008) AGL23, a type I MADS-box gene that controls female gametophyte and embryo development in Arabidopsis. Plant J., 54, 1037–1048.

26. Comai, L., Tyagi, A.P., Winter, K., Holmes-Davis, R., Reynolds, S.H., Stevens, Y. and Byers, B. (2000) Phenotypic instability and rapid gene silencing in newly formed arabidopsis allotetraploids. Plant Cell, 12, 1551–1568.

27. Dart, S., Kron, P. and Mable, B.K. (2004) Characterizing polyploidy inArabidopsis lyratausing chromosome counts and flow cytometry. Can. J. Bot., 82, 185–197.

28. Dolezel, J., Greilhuber, J. and Suda, J. (2007) Estimation of nuclear DNA content in plants using flow cytometry. Nat. Protoc., 2, 2233–2244.

29. Dudchenko, O., Shamim, M.S., Batra, S., et al. (2018) The Juicebox Assembly Tools module facilitates de novo assembly of mammalian genomes with chromosome-length scaffolds for under $1000. bioRxiv, 254797.

30. Durand, N.C., Robinson, J.T., Shamim, M.S., Machol, I., Mesirov, J.P., Lander, E.S. and Aiden, E.L. (2016) Juicebox Provides a Visualization System for Hi-C Contact Maps with Unlimited Zoom. Cell Systems, 3, 99–101.

31. Edgar, R.C. (2004) MUSCLE: multiple sequence alignment with high accuracy and high throughput. Nucleic Acids Res., 32, 1792–1797.

32. Emms, D.M. and Kelly, S. (2019) OrthoFinder: phylogenetic orthology inference for comparative genomics. Genome Biol., 20, 238.

33. English, A.C., Richards, S., Han, Y., et al. (2012) Mind the gap: upgrading genomes with Pacific Biosciences RS long-read sequencing technology. PLoS One, 7, e47768.

34. Erilova, A., Brownfield, L., Exner, V., Rosa, M., Twell, D., Scheid, O.M., Hennig, L. and Köhler, C. (2009) Imprinting of the Polycomb Group Gene MEDEA Serves as a Ploidy Sensor in Arabidopsis. PLoS Genet., 5, e1000663.

35. Florez-Rueda, A.M., Paris, M., Schmidt, A., Widmer, A., Grossniklaus, U. and Städler, T. (2016) Genomic Imprinting in the Endosperm Is Systematically Perturbed in Abortive Hybrid Tomato Seeds. Mol. Biol. Evol., 33, 2935–2946.

36. Flynn, J.M., Hubley, R., Goubert, C., Rosen, J., Clark, A.G., Feschotte, C. and Smit, A.F. (2020) RepeatModeler2 for automated genomic discovery of transposable element families. Proceedings of the National Academy of Sciences, 117, 9451–9457.

37. Folter, S. de, Immink, R.G.H., Kieffer, M., et al. (2005) Comprehensive interaction map of the Arabidopsis MADS Box transcription factors. Plant Cell, 17, 1424–1433.

38. Ghurye, J., Rhie, A., Walenz, B.P., Schmitt, A., Selvaraj, S., Pop, M., Phillippy, A.M. and Koren, S. (2019) Integrating Hi-C links with assembly graphs for chromosome-scale assembly. PLoS Comput. Biol., 15, e1007273.

39. Goel, M., Sun, H., Jiao, W.-B. and Schneeberger, K. (2019) SyRI: finding genomic rearrangements and local sequence differences from whole-genome assemblies. Genome Biol., 20, 277.

40. Gramzow, L. and Theissen, G. (2010) A hitchhiker’s guide to the MADS world of plants. Genome Biol., 11, 214.

41. Gramzow, L. and Theißen, G. (2013) Phylogenomics of MADS-Box Genes in Plants - Two Opposing Life Styles in One Gene Family. Biology, 2, 1150–1164.

42. Gremme, G., Brendel, V., Sparks, M.E. and Kurtz, S. (2005) Engineering a software tool for gene structure prediction in higher organisms. Information and Software Technology, 47, 965–978.

43. Grossniklaus, U., Vielle-Calzada, J.P., Hoeppner, M.A. and Gagliano, W.B. (1998) Maternal control of embryogenesis by MEDEA, a polycomb group gene in Arabidopsis. Science, 280, 446–450.

44. Guan, D., McCarthy, S.A., Wood, J., Howe, K., Wang, Y. and Durbin, R. (2020) Identifying and removing haplotypic duplication in primary genome assemblies. Bioinformatics. Available at: http://dx.doi.org/10.1093/bioinformatics/btaa025 PMID - 31971576.

45. Gurevich, A., Saveliev, V., Vyahhi, N. and Tesler, G. (2013) QUAST: quality assessment tool for genome assemblies. Bioinformatics, 29, 1072–1075.

46. Hao, Z., Lv, D., Ge, Y., Shi, J., Weijers, D., Yu, G. and Chen, J. (2020) RIdeogram: drawing SVG graphics to visualize and map genome-wide data on the idiograms. PeerJ Comput Sci, 6, e251.

47. Hehenberger, E., Kradolfer, D. and Köhler, C. (2012) Endosperm cellularization defines an important developmental transition for embryo development. Development, 139, 2031–2039.

48. Henry, I.M., Dilkes, B.P., Tyagi, A., Gao, J., Christensen, B. and Comai, L. (2014) The BOY NAMED SUE quantitative trait locus confers increased meiotic stability to an adapted natural allopolyploid of Arabidopsis., 26, 181–194.

49. Henschel, K., Kofuji, R., Hasebe, M., Saedler, H., Münster, T. and Theissen, G. (2002) Two ancient classes of MIKC-type MADS-box genes are present in the moss Physcomitrella patens. Mol. Biol. Evol., 19, 801–814.

50. Hoff, K.J., Lange, S., Lomsadze, A., Borodovsky, M. and Stanke, M. (2016) BRAKER1: Unsupervised RNA-Seq-Based Genome Annotation with GeneMark-ET and AUGUSTUS. Bioinformatics, 32, 767–769.

51. Hoff, K.J., Lomsadze, A., Borodovsky, M. and Stanke, M. (2019) Whole-Genome Annotation with BRAKER. In M. Kollmar, ed. Gene Prediction: Methods and Protocols. New York, NY: Springer New York, pp. 65–95.

52. Hohmann, N., Wolf, E.M., Lysak, M.A. and Koch, M.A. (2015) A Time-Calibrated Road Map of Brassicaceae Species Radiation and Evolutionary History. Plant Cell, 27, 2770–2784.

53. Hu, J.-Y., Zhou, Y., He, F., Dong, X., Liu, L.-Y., Coupland, G., Turck, F. and Meaux, J. de (2014) miR824-Regulated AGAMOUS-LIKE16 Contributes to Flowering Time Repression in Arabidopsis. Plant Cell, 26, 2024–2037.

54. Hu, T.T., Pattyn, P., Bakker, E.G., et al. (2011) The Arabidopsis lyrata genome sequence and the basis of rapid genome size change. Nat. Genet., 43, 476–481.

55. Jaegle, B., Pisupati, R., Soto-Jiménez, L.M., Burns, R., Rabanal, F.A. and Nordborg, M. (2023) Extensive sequence duplication in Arabidopsis revealed by pseudo-heterozygosity. Genome Biol., 24, 44.

56. Jiang, X., Song, Q., Ye, W. and Chen, Z.J. (2021) Concerted genomic and epigenomic changes accompany stabilization of Arabidopsis allopolyploids. Nature Ecology & Evolution, 1–12.

57. Jin, J.-J., Yu, W.-B., Yang, J.-B., Song, Y., dePamphilis, C.W., Yi, T.-S. and Li, D.-Z. (2020) GetOrganelle: a fast and versatile toolkit for accurate de novo assembly of organelle genomes. Genome Biol., 21, 241.

58. Johnston, J.S., Pepper, A.E., Hall, A.E., Chen, Z.J., Hodnett, G., Drabek, J., Lopez, R. and Price, H.J. (2005) Evolution of genome size in Brassicaceae. Ann. Bot., 95, 229–235.

59. Jørgensen, M.H., Ehrich, D., Schmickl, R., Koch, M.A. and Brysting, A.K. (2011) Interspecific and interploidal gene flow in Central European Arabidopsis (Brassicaceae). BMC Evol. Biol., 11, 346.

60. Josefsson, C., Dilkes, B. and Comai, L. (2006) Parent-dependent loss of gene silencing during interspecies hybridization. Curr. Biol., 16, 1322–1328.

61. Kang, I.-H., Steffen, J.G., Portereiko, M.F., Lloyd, A. and Drews, G.N. (2008) The AGL62 MADS domain protein regulates cellularization during endosperm development in Arabidopsis. Plant Cell, 20, 635–647.

62. Katoh, K. and Standley, D.M. (2013) MAFFT multiple sequence alignment software version 7: improvements in performance and usability. Mol. Biol. Evol., 30, 772–780.

63. Kaufmann, K., Muiño, J.M., Jauregui, R., Airoldi, C.A., Smaczniak, C., Krajewski, P. and Angenent, G.C. (2009) Target genes of the MADS transcription factor SEPALLATA3: integration of developmental and hormonal pathways in the Arabidopsis flower. PLoS Biol., 7, e1000090.

64. Kawabe, A. and Charlesworth, D. (2006) Patterns of DNA Variation Among Three Centromere Satellite Families in Arabidopsis halleri and A. lyrata. J. Mol. Evol., 64, 237.

65. Kemi, U., Niittyvuopio, A., Toivainen, T., Pasanen, A., Quilot-Turion, B., Holm, K., Lagercrantz, U., Savolainen, O. and Kuittinen, H. (2013) Role of vernalization and of duplicated FLOWERING LOCUS C in the perennial Arabidopsis lyrata. New Phytologist, 197, 323–335. Available at: http://dx.doi.org/10.1111/j.1469-8137.2012.04378.x.

66. Kim, D., Paggi, J.M., Park, C., Bennett, C. and Salzberg, S.L. (2019) Graph-based genome alignment and genotyping with HISAT2 and HISAT-genotype. Nat. Biotechnol., 37, 907–915.

67. Kinoshita, T., Yadegari, R., Harada, J.J., Goldberg, R.B. and Fischer, R.L. (1999) Imprinting of the MEDEA polycomb gene in the Arabidopsis endosperm. Plant Cell, 11, 1945–1952.

68. Kirkbride, R.C., Lu, J., Zhang, C., Mosher, R.A., Baulcombe, D.C. and Chen, Z.J. (2019) Maternal small RNAs mediate spatial-temporal regulation of gene expression, imprinting, and seed development in Arabidopsis. Proc. Natl. Acad. Sci. U. S. A., 201807621.

69. Klosinska, M., Picard, C.L. and Gehring, M. (2016) Conserved imprinting associated with unique epigenetic signatures in the Arabidopsis genus. Nature Plants, 2, 16145.

70. Köhler, C., Hennig, L., Spillane, C., Pien, S., Gruissem, W. and Grossniklaus, U. (2003) The Polycomb-group protein MEDEA regulates seed development by controlling expression of the MADS-box gene PHERES1. Genes Dev., 17, 1540–1553.

71. Kolesnikova, U.K., Scott, A.D., Van de Velde, J.D., et al. (2023) Transition to self-compatibility associated with dominant S-allele in a diploid Siberian progenitor of allotetraploid Arabidopsis kamchatica revealed by Arabidopsis lyrata genomes. bioRxiv, 2022.06.24.497443. Available at: https://www.biorxiv.org/content/10.1101/2022.06.24.497443v2 [Accessed May 18, 2023].

72. Kolmogorov, M., Yuan, J., Lin, Y. and Pevzner, P.A. (2019) Assembly of long, error-prone reads using repeat graphs. Nat. Biotechnol., 37, 540–546.

73. Koren, S., Walenz, B.P., Berlin, K., Miller, J.R., Bergman, N.H. and Phillippy, A.M. (2017) Canu: scalable and accurate long-read assembly via adaptive k-mer weighting and repeat separation. Genome Res., 27, 722–736.

74. Kovaka, S., Ou, S., Jenike, K.M. and Schatz, M.C. (2023) Approaching complete genomes, transcriptomes and epi-omes with accurate long-read sequencing. Nat. Methods, 20, 12–16.

75. Kress, W.J., Soltis, D.E., Kersey, P.J., Wegrzyn, J.L., Leebens-Mack, J.H., Gostel, M.R., Liu, X. and Soltis, P.S. (2022) Green plant genomes: What we know in an era of rapidly expanding opportunities. Proc. Natl. Acad. Sci. U. S. A., 119. Available at: http://dx.doi.org/10.1073/pnas.2115640118.

76. Kronenberg, Z.N., Rhie, A., Koren, S., et al. (2021) Extended haplotype-phasing of long-read de novo genome assemblies using Hi-C. Nat. Commun., 12, 1935.

77. Krueger, F. (2016) TrimGalore: A wrapper around Cutadapt and FastQC to consistently apply adapter and quality trimming to FastQ files, with extra functionality for RRBS data. TrimGalore (accessed on 27 August 2019).

78. Lafon-Placette, C., Johannessen, I.M., Hornslien, K.S., et al. (2017) Endosperm-based hybridization barriers explain the pattern of gene flow between Arabidopsis lyrata and Arabidopsis arenosa in Central Europe. Proc. Natl. Acad. Sci. U. S. A., 114, E1027–E1035.

79. Legrand, S., Caron, T., Maumus, F., et al. (2019) Differential retention of transposable element-derived sequences in outcrossing Arabidopsis genomes. Mob. DNA, 10, 30.

80. Leinonen, R., Sugawara, H., Shumway, M. and on behalf of the International Nucleotide Sequence Database Collaboration (2011) The sequence read archive. Nucleic Acids Res., 39, D19–D21. Available at: [Accessed January 20, 2023].

81. Liao, Y., Smyth, G.K. and Shi, W. (2014) featureCounts: an efficient general purpose program for assigning sequence reads to genomic features. Bioinformatics, 30, 923–930.

82. Li, H. (2013) Aligning sequence reads, clone sequences and assembly contigs with BWA-MEM. arXiv [q-bio.GN]. Available at: http://arxiv.org/abs/1303.3997.

83. Li, H. (2018) Minimap2: pairwise alignment for nucleotide sequences. Bioinformatics, 34, 3094–3100.

84. Li, H. and Durbin, R. (2009) Fast and accurate short read alignment with Burrows-Wheeler transform. Bioinformatics, 25, 1754–1760.

85. Lindsey, B.E., 3rd, Rivero, L., Calhoun, C.S., Grotewold, E. and Brkljacic, J. (2017) Standardized Method for High-throughput Sterilization of Arabidopsis Seeds. J. Vis. Exp. Available at: http://dx.doi.org/10.3791/56587.

86. Liu, Z., Cheema, J., Vigouroux, M., Hill, L., Reed, J., Paajanen, P., Yant, L. and Osbourn, A. (2020) Formation and diversification of a paradigm biosynthetic gene cluster in plants. Nat. Commun., 11, 5354.

87. Love, M.I., Huber, W. and Anders, S. (2014) Moderated estimation of fold change and dispersion for RNA-seq data with DESeq2. Genome Biol., 15, 550.

88. Lysak, M.A., Berr, A., Pecinka, A., Schmidt, R., McBreen, K. and Schubert, I. (2006) Mechanisms of chromosome number reduction in Arabidopsis thaliana and related Brassicaceae species. Proc. Natl. Acad. Sci. U. S. A., 103, 5224–5229.

89. Lysak, M.A., Koch, M.A., Beaulieu, J.M., Meister, A. and Leitch, I.J. (2009) The dynamic ups and downs of genome size evolution in Brassicaceae. Mol. Biol. Evol., 26, 85–98.

90. Maheshwari, S., Ishii, T., Brown, C.T., Houben, A. and Comai, L. (2017) Centromere location in Arabidopsis is unaltered by extreme divergence in CENH3 protein sequence. Genome Res., 27, 471–478.

91. Makarevich, G., Villar, C.B.R., Erilova, A. and Köhler, C. (2008) Mechanism of PHERES1 imprinting in Arabidopsis. J. Cell Sci., 121, 906–912.

92. Manni, M., Berkeley, M.R., Seppey, M., Simão, F.A. and Zdobnov, E.M. (2021) BUSCO Update: Novel and Streamlined Workflows along with Broader and Deeper Phylogenetic Coverage for Scoring of Eukaryotic, Prokaryotic, and Viral Genomes. Mol. Biol. Evol., 38, 4647–4654.

93. Marburger, S., Monnahan, P., Seear, P.J., et al. (2019) Interspecific introgression mediates adaptation to whole genome duplication. Nat. Commun., 10, 5218.

94. Masiero, S., Colombo, L., Grini, P.E., Schnittger, A. and Kater, M.M. (2011) The emerging importance of type I MADS box transcription factors for plant reproduction. Plant Cell, 23, 865– 872.

95. Minh, B.Q., Schmidt, H.A., Chernomor, O., Schrempf, D., Woodhams, M.D., Haeseler, A. von and Lanfear, R. (2020) IQ-TREE 2: New models and efficient methods for phylogenetic inference in the genomic era. Mol. Biol. Evol. Available at: http://dx.doi.org/10.1093/molbev/msaa015.

96. Müller-Xing, R., Ardiansyah, R., Xing, Q., et al. (2022) Polycomb proteins control floral determinacy by H3K27me3-mediated repression of pluripotency genes in Arabidopsis thaliana. J. Exp. Bot., 73, 2385–2402.

97. Murashige, T. and Skoog, F. (1962) A revised medium for rapid growth and bio assays with tobacco tissue cultures. Physiol. Plant., 15, 473–497.

98. Naish, M., Alonge, M., Wlodzimierz, P., et al. (2021) The genetic and epigenetic landscape of the Arabidopsis centromeres. Science, 374, eabi7489.

99. Nam, J., Kim, J., Lee, S., An, G., Ma, H. and Nei, M. (2004) Type I MADS-box genes have experienced faster birth-and-death evolution than type II MADS-box genes in angiosperms. Proc. Natl. Acad. Sci. U. S. A., 101, 1910–1915.

100. Nasrallah, M.E., Yogeeswaran, K., Snyder, S. and Nasrallah, J.B. (2000) Arabidopsis species hybrids in the study of species differences and evolution of amphiploidy in plants. Plant Physiol., 124, 1605–1614.

101. Nattestad, M. and Schatz, M.C. (2016) Assemblytics: a web analytics tool for the detection of variants from an assembly. Bioinformatics, 32, 3021–3023.

102. Novikova, P.Y., Hohmann, N., Nizhynska, V., et al. (2016) Sequencing of the genus Arabidopsis identifies a complex history of nonbifurcating speciation and abundant trans-specific polymorphism. Nature Publishing Group. Available at: http://dx.doi.org/10.1038/ng.3617.

103. Novikova, P.Y., Tsuchimatsu, T., Simon, S., et al. (2017) Genome sequencing reveals the origin of the allotetraploid Arabidopsis suecica. *Mol. Biol. Evol.*, msw299.

104. N Schep, A. and K Kummerfeld, S. (2017) iheatmapr: Interactive complex heatmaps in R. J. Open Source Softw., 2, 359.

105. Paape, T., Briskine, R.V., Halstead-Nussloch, G., et al. (2018) Patterns of polymorphism and selection in the subgenomes of the allopolyploid Arabidopsis kamchatica. Nat. Commun., 9, 3909.

106. Parenicová, L., Folter, S. de, Kieffer, M., et al. (2003) Molecular and phylogenetic analyses of the complete MADS-box transcription factor family in Arabidopsis: new openings to the MADS world. Plant Cell, 15, 1538–1551.

107. Pelaz, S., Tapia-López, R., Alvarez-Buylla, E.R. and Yanofsky, M.F. (2001) Conversion of leaves into petals in Arabidopsis. Curr. Biol., 11, 182–184.

108. Pellicer, J. and Leitch, I.J. (2020) The Plant DNA C-values database (release 7.1): an updated online repository of plant genome size data for comparative studies. New Phytol., 226, 301–305.

109. Price, M.N., Dehal, P.S. and Arkin, A.P. (2010) FastTree 2--approximately maximum-likelihood trees for large alignments. PLoS One, 5, e9490.

110. Pucker, B., Irisarri, I., Vries, J. de and Xu, B. (2022) Plant genome sequence assembly in the era of long reads: Progress, challenges and future directions. Quantitative Plant Biology, 3, e5. Available at: [Accessed November 11, 2022].

111. Qiu, Y. and Köhler, C. (2022) Endosperm Evolution by Duplicated and Neofunctionalized Type I MADS-Box Transcription Factors. Mol. Biol. Evol., 39. Available at: http://dx.doi.org/10.1093/molbev/msab355.

112. Qiu, Y. and Köhler, C. (2020) Mobility connects: transposable elements wire new transcriptional networks by transferring transcription factor binding motifs. Biochem. Soc. Trans. Available at: http://dx.doi.org/10.1042/bst20190937.

113. Rachael Workman, Winston Timp, Renee Fedak, Duncan Kilburn, Stephanie Hao and Kelvin Liu (2021) High Molecular Weight DNA Extraction from Recalcitrant Plant Species for Third Generation Sequencing. Protocol Exchange. Available at: https://doi.org/10.1038/protex.2018.059.

114. Ranallo-Benavidez, T.R., Jaron, K.S. and Schatz, M.C. (2020) GenomeScope 2.0 and Smudgeplot for reference-free profiling of polyploid genomes. Nat. Commun., 11, 1432.

115. Ratcliffe, O.J., Kumimoto, R.W., Wong, B.J. and Riechmann, J.L. (2003) Analysis of the Arabidopsis MADS AFFECTING FLOWERING Gene Family: MAF2 Prevents Vernalization by Short Periods of Cold. The Plant Cell Online, 15, 1159–1169.

116. Rawat, V., Abdelsamad, A., Pietzenuk, B., Seymour, D.K., Koenig, D., Weigel, D., Pecinka, A. and Schneeberger, K. (2015) Improving the Annotation of Arabidopsis lyrata Using RNA-Seq Data. PLoS One, 10, e0137391.

117. Rebernig, C.A., Lafon-Placette, C., Hatorangan, M.R., Slotte, T. and Köhler, C. (2015) Non-reciprocal Interspecies Hybridization Barriers in the Capsella Genus Are Established in the Endosperm. PLoS Genet., 11, e1005295.

118. Rhie, A., McCarthy, S.A., Fedrigo, O., et al. (2021) Towards complete and error-free genome assemblies of all vertebrate species. Nature, 592, 737–746.

119. Rhie, A., Walenz, B.P., Koren, S. and Phillippy, A.M. (2020) Merqury: reference-free quality, completeness, and phasing assessment for genome assemblies. Genome Biol., 21, 2020.03.15.992941.

120. Roach, M.J., Schmidt, S.A. and Borneman, A.R. (2018) Purge Haplotigs: allelic contig reassignment for third-gen diploid genome assemblies. BMC Bioinformatics, 19, 460.

121. Robinson, J.T., Thorvaldsdóttir, H., Winckler, W., Guttman, M., Lander, E.S., Getz, G. and Mesirov, J.P. (2011) Integrative genomics viewer. Nat. Biotechnol., 29, 24–26.

122. Roszak, P. and Köhler, C. (2011) Polycomb group proteins are required to couple seed coat initiation to fertilization. Proceedings of the National Academy of Sciences, 108, 20826–20831.

123. Salmela, L. and Rivals, E. (2014) LoRDEC: accurate and efficient long read error correction. Bioinformatics, 30, 3506–3514.

124. Schaffer, R.J., Ireland, H.S., Ross, J.J., Ling, T.J. and David, K.M. (2013) SEPALLATA1/2- suppressed mature apples have low ethylene, high auxin and reduced transcription of ripening-related genes. AoB Plants, 5, ls047.

125. Schmickl, R. and Koch, M.A. (2011) Arabidopsis hybrid speciation processes. Proc. Natl. Acad. Sci. U. S. A., 108, 14192–14197.

126. Schmickl, R. and Yant, L. (2021) Adaptive introgression: how polyploidy reshapes gene flow landscapes. New Phytol., 230, 457–461.

127. Schönrock, N., Bouveret, R., Leroy, O., Borghi, L., Köhler, C., Gruissem, W. and Hennig, L. (2006) Polycomb-group proteins repress the floral activator AGL19 in the FLC-independent vernalization pathway. Genes Dev., 20, 1667–1678.

128. Seymour, G.B., Ryder, C.D., Cevik, V., Hammond, J.P., Popovich, A., King, G.J., Vrebalov, J., Giovannoni, J.J. and Manning, K. (2011) A SEPALLATA gene is involved in the development and ripening of strawberry (Fragaria x ananassa Duch.) fruit, a non-climacteric tissue. J. Exp. Bot., 62, 1179–1188.

129. Shirzadi, R., Andersen, E.D., Bjerkan, K.N., et al. (2011) Genome-Wide Transcript Profiling of Endosperm without Paternal Contribution Identifies Parent-of-Origin–Dependent Regulation of AGAMOUS-LIKE36. PLoS Genetics, 7, e1001303. Available at: http://dx.doi.org/10.1371/journal.pgen.1001303.

130. Simão, F.A., Waterhouse, R.M., Ioannidis, P., Kriventseva, E.V. and Zdobnov, E.M. (2015) BUSCO: assessing genome assembly and annotation completeness with single-copy orthologs. Bioinformatics, 31, 3210–3212.

131. Slotte, T., Hazzouri, K.M., Ågren, J.A., et al. (2013) The Capsella rubella genome and the genomic consequences of rapid mating system evolution. Nat. Genet., 45, 831–835.

132. Smit, A.F.A., Hubley, R. and Green, P. (2015) RepeatMasker Open-4.0. 2013--2015.

133. Soppe, W.J.J., Viñegra de la Torre, N. and Albani, M.C. (2021) The Diverse Roles of FLOWERING LOCUS C in Annual and Perennial Brassicaceae Species. Front. Plant Sci., 12, 627258.

134. Steffen, J.G., Kang, I.-H., Portereiko, M.F., Lloyd, A. and Drews, G.N. (2008) AGL61 interacts with AGL80 and is required for central cell development in Arabidopsis. Plant Physiol., 148, 259–268.

135. Storer, J., Hubley, R., Rosen, J., Wheeler, T.J. and Smit, A.F. (2021) The Dfam community resource of transposable element families, sequence models, and genome annotations. Mob. DNA, 12, 2.

136. Thangavel, G. and Nayar, S. (2018) A Survey of MIKC Type MADS-Box Genes in Non-seed Plants: Algae, Bryophytes, Lycophytes and Ferns. Front. Plant Sci., 9, 510.

137. The Arabidopsis Genome Initiative (2000) Analysis of the genome sequence of the flowering plant Arabidopsis thaliana. Nature, 408, 796–815. Available at: http://dx.doi.org/10.1038/35048692.

138. Tonosaki, K., Sekine, D., Ohnishi, T., Ono, A., Furuumi, H., Kurata, N. and Kinoshita, T. (2017) Overcoming the species hybridization barrier by ploidy manipulation in the genus Oryza. Plant J. Available at: http://dx.doi.org/10.1111/tpj.13803.

139. Verelst, W., Saedler, H. and Münster, T. (2007) MIKC* MADS-protein complexes bind motifs enriched in the proximal region of late pollen-specific Arabidopsis promoters. Plant Physiol., 143, 447–460.

140. Walia, H., Josefsson, C., Dilkes, B., Kirkbride, R., Harada, J. and Comai, L. (2009) Dosage-dependent deregulation of an AGAMOUS-LIKE gene cluster contributes to interspecific incompatibility. Curr. Biol., 19, 1128–1132.

141. Walker, B.J., Abeel, T., Shea, T., et al. (2014) Pilon: an integrated tool for comprehensive microbial variant detection and genome assembly improvement. PLoS One, 9, e112963.

142. Wang, Y., Tang, H., Debarry, J.D., et al. (2012) MCScanX: a toolkit for detection and evolutionary analysis of gene synteny and collinearity. Nucleic Acids Res., 40, e49.

143. Waterhouse, R.M., Seppey, M., Simão, F.A., Manni, M., Ioannidis, P., Klioutchnikov, G., Kriventseva, E.V. and Zdobnov, E.M. (2018) BUSCO Applications from Quality Assessments to Gene Prediction and Phylogenomics. Mol. Biol. Evol., 35, 543–548.

144. Yant, L., Hollister, J.D., Wright, K.M., Arnold, B.J., Higgins, J.D., Franklin, F.C.H. and Bomblies, K. (2013) Meiotic adaptation to genome duplication in Arabidopsis arenosa. Curr. Biol., 23, 2151–2156.

145. Yoshida, T. and Kawabe, A. (2013) Importance of gene duplication in the evolution of genomic imprinting revealed by molecular evolutionary analysis of the type I MADS-box gene family in Arabidopsis species. PLoS One, 8, e73588.

146. Zhang, S., Wang, D., Zhang, H., et al. (2018) FERTILIZATION-INDEPENDENT SEED-Polycomb Repressive Complex 2 plays a dual role in regulating type I MADS-box genes in early endosperm development. Plant Physiol., 00534.2017.

147. Zhang, X., Zhang, S., Zhao, Q., Ming, R. and Tang, H. (2019) Assembly of allele-aware, chromosomal-scale autopolyploid genomes based on Hi-C data. Nature Plants, 5, 833–845.

148. Zimin, A.V., Marçais, G., Puiu, D., Roberts, M., Salzberg, S.L. and Yorke, J.A. (2013) The MaSuRCA genome assembler. Bioinformatics, 29, 2669–2677.

