## Supplementary Figure S1: Pipeline for genome assembly for "Structural evidence for MADS-box type I family expansion seen in new assemblies of *A. arenosa* and *A. lyrata*"

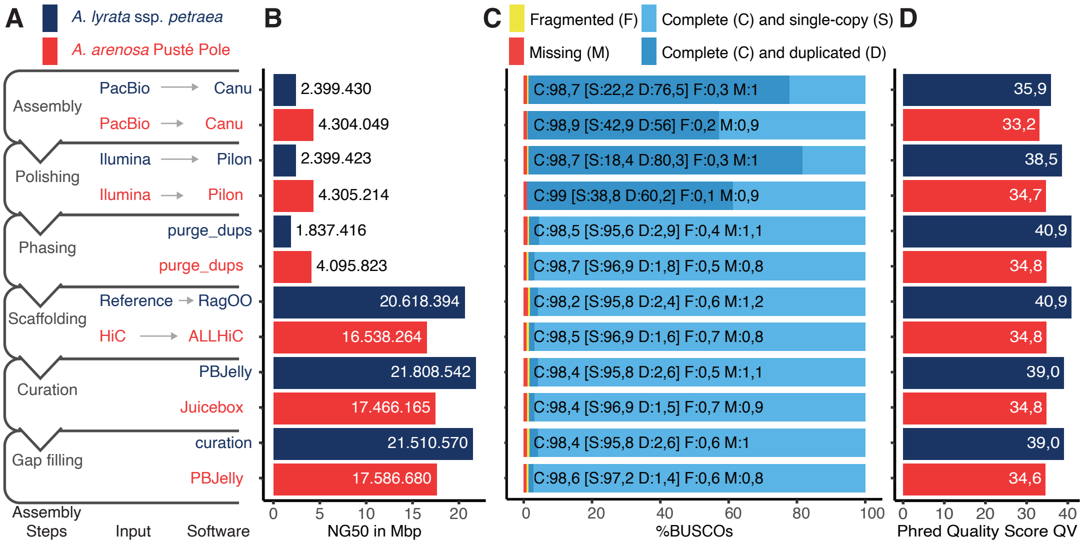


**Figure S1: Pipeline for genome assembly.** *A. lyrata* ssp. *petraea* (blue) and *A. arenosa* Pusté Pole (red). *A. arenosa* was assembled de novo from Pacbio long-read data and scaffolded using Hi-C data. For *A. lyrata* ssp. *petraea* a draft assembly was made de novo from PacBio long-reads, which were subsequently scaffolded against the published genome of *A. lyrata* ssp. *lyrata* (Hu et al., 2011). (A) Pipeline indicating each step in the assembly process. For each step additional programs or methods were tested; details of these analyses can be found in Data S1. (B) Statistics used in quality control and software decisions. Sequence contiguity is shown in NG50, using the calculated genome size of 201,144,702 bp for *A. lyrata* ssp. *petraea*, and 179,232,250 bp for *A. arenosa* Pusté Pole. (C) BUSCO scores using the gene set for Embryophyta odb9 with 1440 single-copy genes. For the assembly steps, completeness and duplication can be seen in light and dark blue, respectively. The combination of PacBio long reads with Canu created near-complete assemblies. Allelic duplications were identified and removed using Purge_Dups. (D) Phred Quality Score (QV) was calculated with Merqury (Rhie et al., 2020). A QV of 30 corresponds to 99.9% accuracy and QV 40 to 99.99%. The quality of both long read assemblies was improved by the Pilon polishing step using Illumina reads. While the *A. lyrata* ssp. *petraea* assembly shows a consistently lower error rate, both assemblies have high quality during all assembly steps. For more assemblies and detailed statistics see Data S1.
