## Supplementary Figure S2: GenomeScope profiles for "Structural evidence for MADS-box type I family expansion seen in new assemblies of *A. arenosa* and *A. lyrata*"

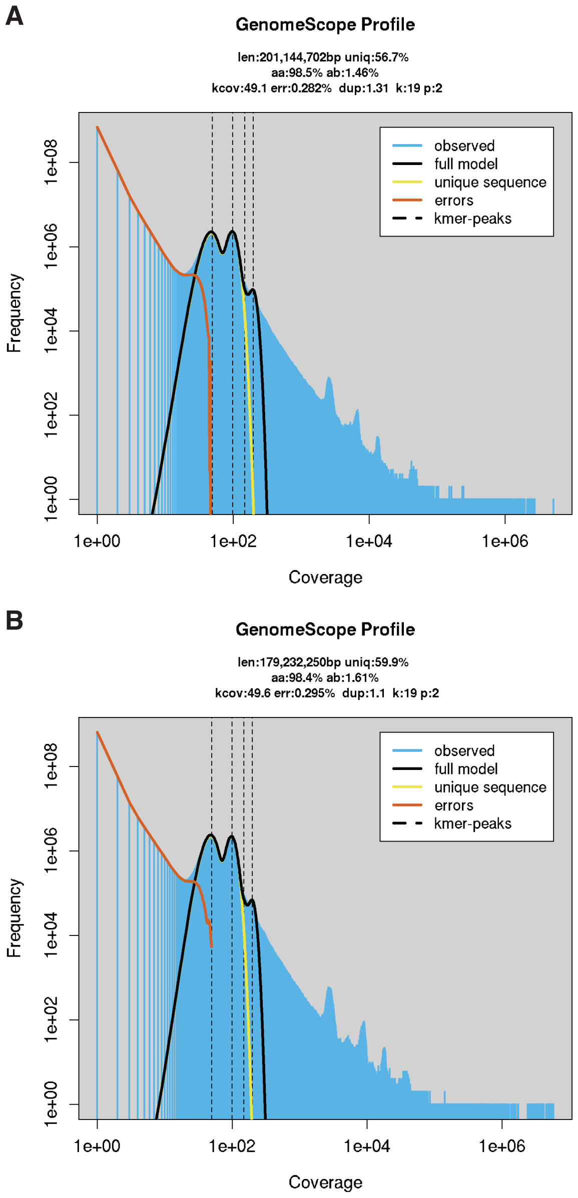


**Figure S2: GenomeScope profiles.** Mercury produced the k-mer frequency spectra of Illumina reads (blue) for a k-mer length of 19 nt. To this spectrum, the GenomeScope model (black line) was fitted, and genome length, repetitiveness, and heterozygosity rates were predicted. (A) K-mer spectrum for *A. lyrata* ssp*. petraea.* GenomeScope v2.0 infers a genome length of 201,144,702 bp with a heterozygosity rate of 1.46%. The heterozygous and homozygous peaks were detected with an approximal coverage of 50 and 100, respectively. K-mers with coverage above 1000 were enriched for ribosomal, mitochondrial, and chloroplast DNA. (B) K-mer spectrum for *A. arenosa* Psuté Pole. The heterozygous and homozygous peaks were detected with an approximal coverage of 50 and 100, respectively. GenomeScope infers a genome length of 179,232,250 bp with a heterozygosity rate of 1.61%.
