## Supplementary Figure S3: Copy number spectrum plots for "Structural evidence for MADS-box type I family expansion seen in new assemblies of *A. arenosa* and *A. lyrata*"

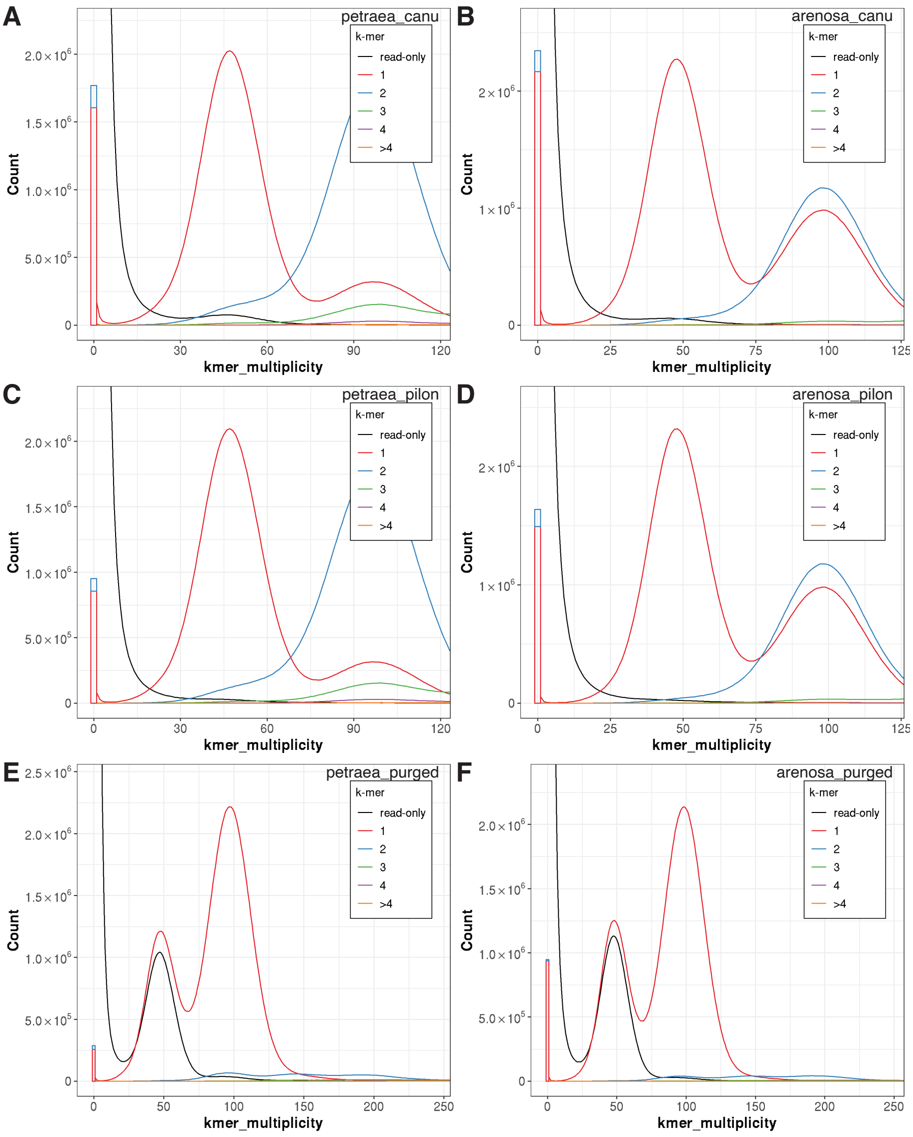


**Figure S3: Copy number spectrum plots.** For quality control, the k-mer frequencies from every assembly were compared to the k-mer frequencies of the corresponding Illumina reads. The k-mer spectrum shows how many unique 19k-mer exist (Count) with a specific coverage (k-mer multiplicity). Each Illumina read k-mer is further grouped and colored depending on how often it is found in the assemblies. The first peak at half coverage is expected to contain k-mers found only on one haplotype, while the second peak should include k-mer from both haplotypes. (A) Copy number spectrum plot (spectra-cn) for the primary *A. lyrata* ssp. *petraea* assembly. The assembly was created with the Canu assembler using PacBio long reads. The spectrum indicates a high-quality assembly where nearly all k-mers found in the Illumina reads (not used for the assembly) were also detected in the assembly with their expected frequencies. K-mers found only in the long-read assembly are displayed as a red/blue-bar to the left of the spectrum plot. (B) Spectra-cn plot for the primary *A. arenosa* Pusté Pole assembly. The Canu assembler using uncorrected PacBio long-reads created the most complete draft assembly. Slightly reduced haploid resolution compared to the *A. lyrata* ssp. *petraea* assembly. (C) Spectra-cn plot of the *A. lyrata* ssp. *petraea* assembly after the Pilon polishing step. Using Illumina reads for polishing reduced the number of suspected erroneous k-mers found only in the assembly (blue/red bar on the left). (D) Spectra-cn plot of the *A. arenosa* Pusté Pole assembly after the Pilon polishing step. (E) Spectra-cn plot of the *A. lyrata* ssp. *petraea* assembly after the haplotic purging step. Only the primary haplotig is compared to the Illumina read set. (F) Spectra-cn plot of the *A. arenosa* Pusté Pole assembly after the haplotig purging. K-mer frequencies for the primary haplotig are shown. Half of the single-copy k-mers are missing and are found in the alternative haplotig contigs.
