## Supplementary Figure S4: A. lyrata ssp. petraea - A. lyrata ssp. lyrata alignments for "Structural evidence for MADS-box type I family expansion seen in new assemblies of *A. arenosa* and *A. lyrata*"

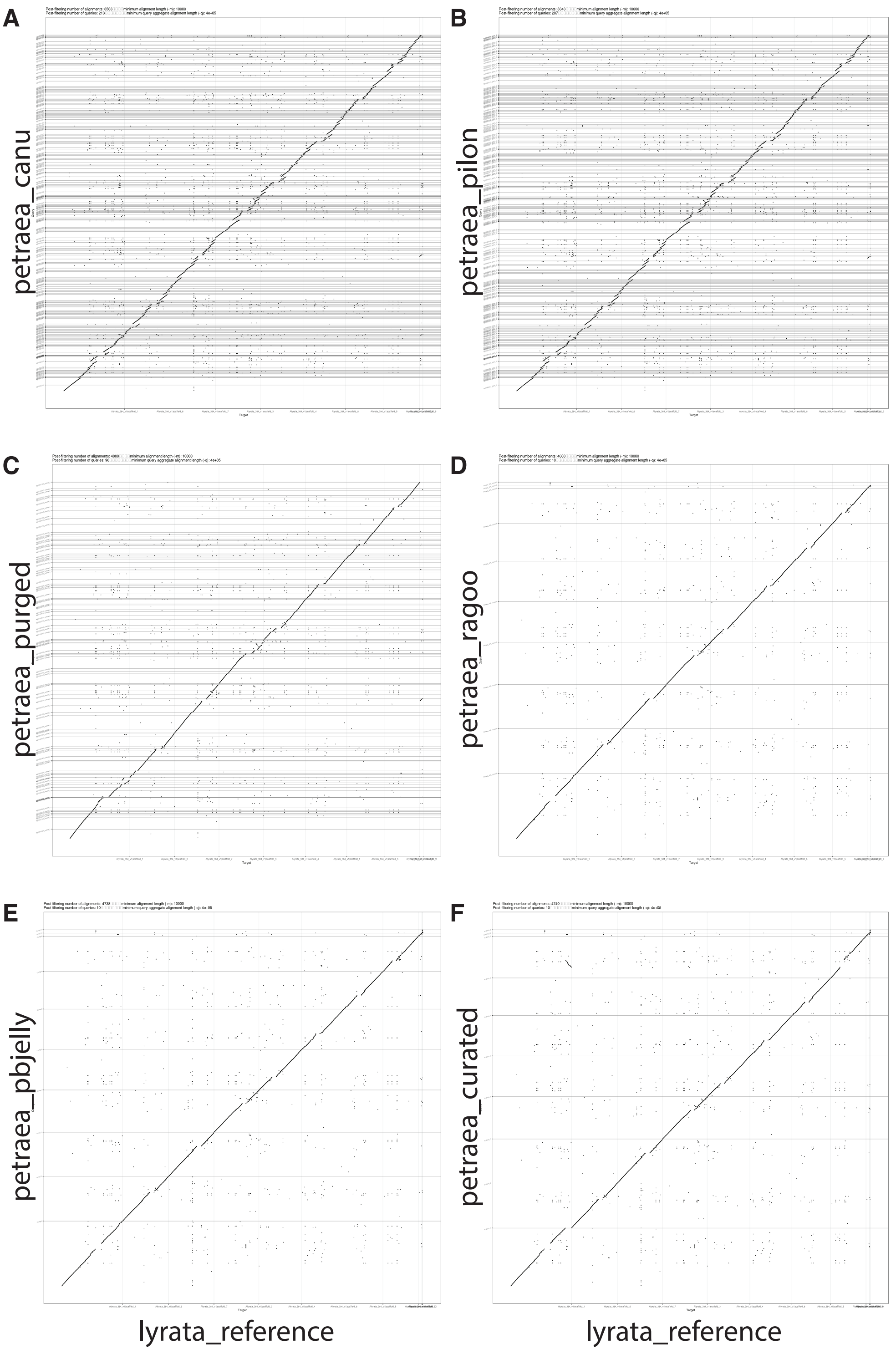


**Figure S4: *A. lyrata* ssp. *petraea*-*A. lyrata* ssp. *lyrata* alignments.** After each processing step, all *A. lyrata* ssp. *petraea* contigs and scaffolds were aligned to the *A. lyrata* ssp. *lyrata* reference genome as quality control. Larger mis-assemblies were easily detected and excluded. The reference scaffolds are on the x-axis, while the respective new assemblies are sorted on the y-axis. (A) Alignment of the *A. lyrata* ssp. *petraea* Canu draft assembly. (B) Alignment of the contigs after the Pilon polishing step shows a high duplication rate compared to the haploid reference. (C) Alignment of the contigs after the haplotic purging step. Contiguity remained after the removal of duplicated sequences. (D) Reference alignment after the reference-based scaffolding step using RaGoo. (E) Alignment after the gab closure by PBJelly displays no considerable changes. (F) Alignment of our final curated *A. lyrata* ssp. *petraea* assembly. The rearrange-ment of the previously reported misassembly for *A. lyrata* ssp. *lyrata* can be seen between scaffolds one and two (sorted by size) in the top left corner.
