## Supplemental Data 1 for "Structural evidence for MADS-box type I family expansion seen in new assemblies of *A. arenosa* and *A. lyrata*"

Supplementary Figure S5

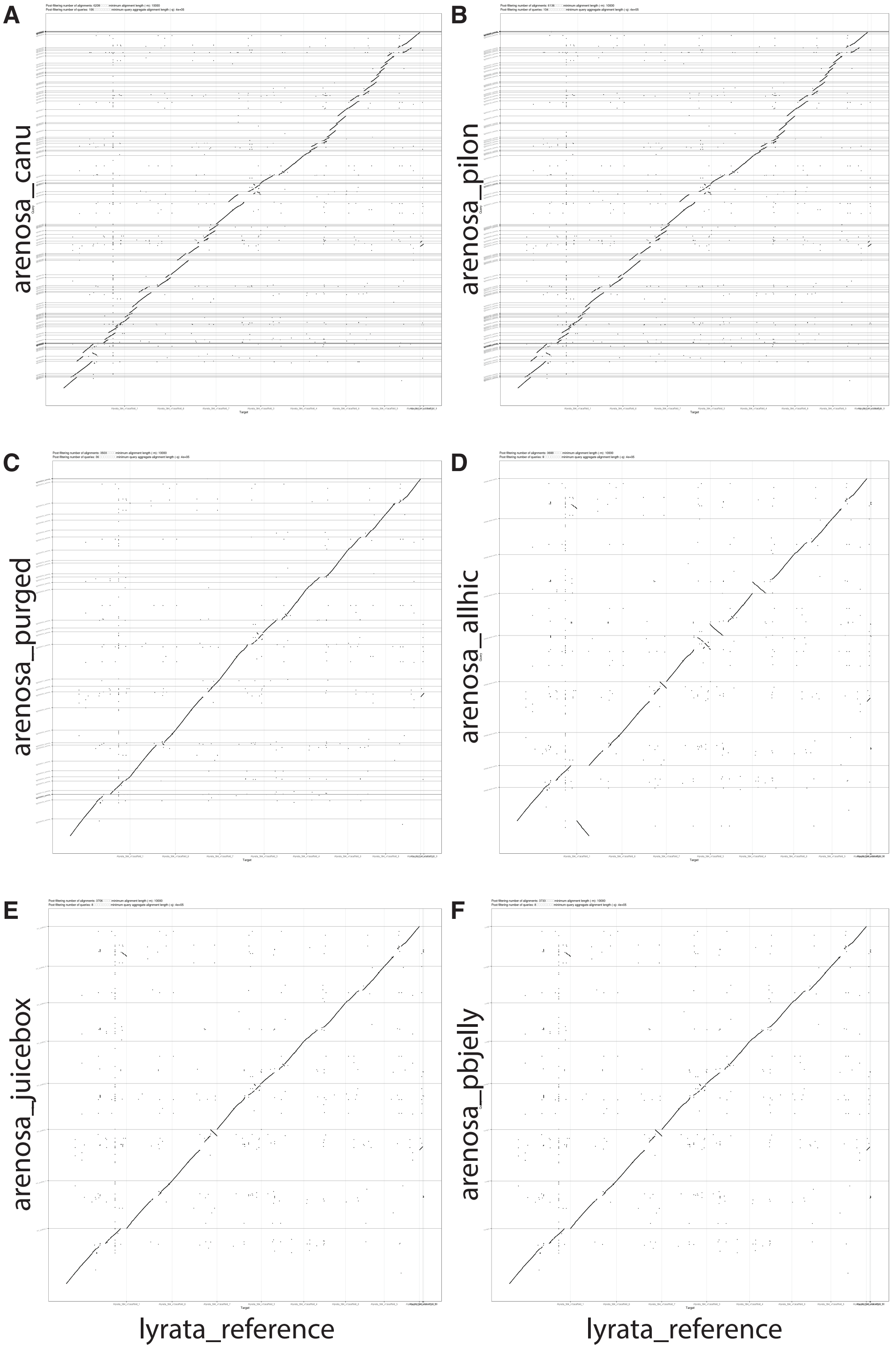


**Figure S5:** ***A. arenosa*** **Pusté Pole-*A. lyrata*** **ssp.** ***lyrata*** **alignments.** For quality control, all *A. arenosa* Pusté Pole sequences were aligned to the *A. lyrata* ssp. *lyrata* reference genome. Although structural variation is expected between the species, inconsistent rearrangements between assemblies were fast detected and excluded. The y-axis represents the length sorted scaffolds from the published *A. lyrata* ssp. *lyrata* genome. (A) Contigs of the *A. arenosa* Pusté Pole Canu draft assembly on the y-axis show a high number of overlaps. (B) Alignment of the same contigs after the Pilon polishing step. (C) Alignment of the contigs after the haplotic purging step. Only the primary haplotig is shown. Contiguity remains after the removal of duplicated sequences. (D) The alignment plots were especially helpful to remove erroneous Hi-C scaffolding approaches. The best performance displayed here is our selected approach using ALLHiC. (E) Minor scaffolding mistakes were curated with the Juicebox Assembly Tools (JBAT) based on the Hi-C linkage map. The curated result aligned astoundingly well to the *A. lyrata* ssp. *lyrata* scaffolds. (F) Gap-closure with PBJelly did not change the scaffold arrangement. The previously reported misassembly on *A. lyrata* ssp. *lyrata* scaffold can be seen in the top left corner. In addition, one more extensive inversion is displayed on scaffold 7, in the center of the plot.
