## Supplemental Data 2 for "Structural evidence for MADS-box type I family expansion seen in new assemblies of *A. arenosa* and *A. lyrata*"

Supplementary Figure S6

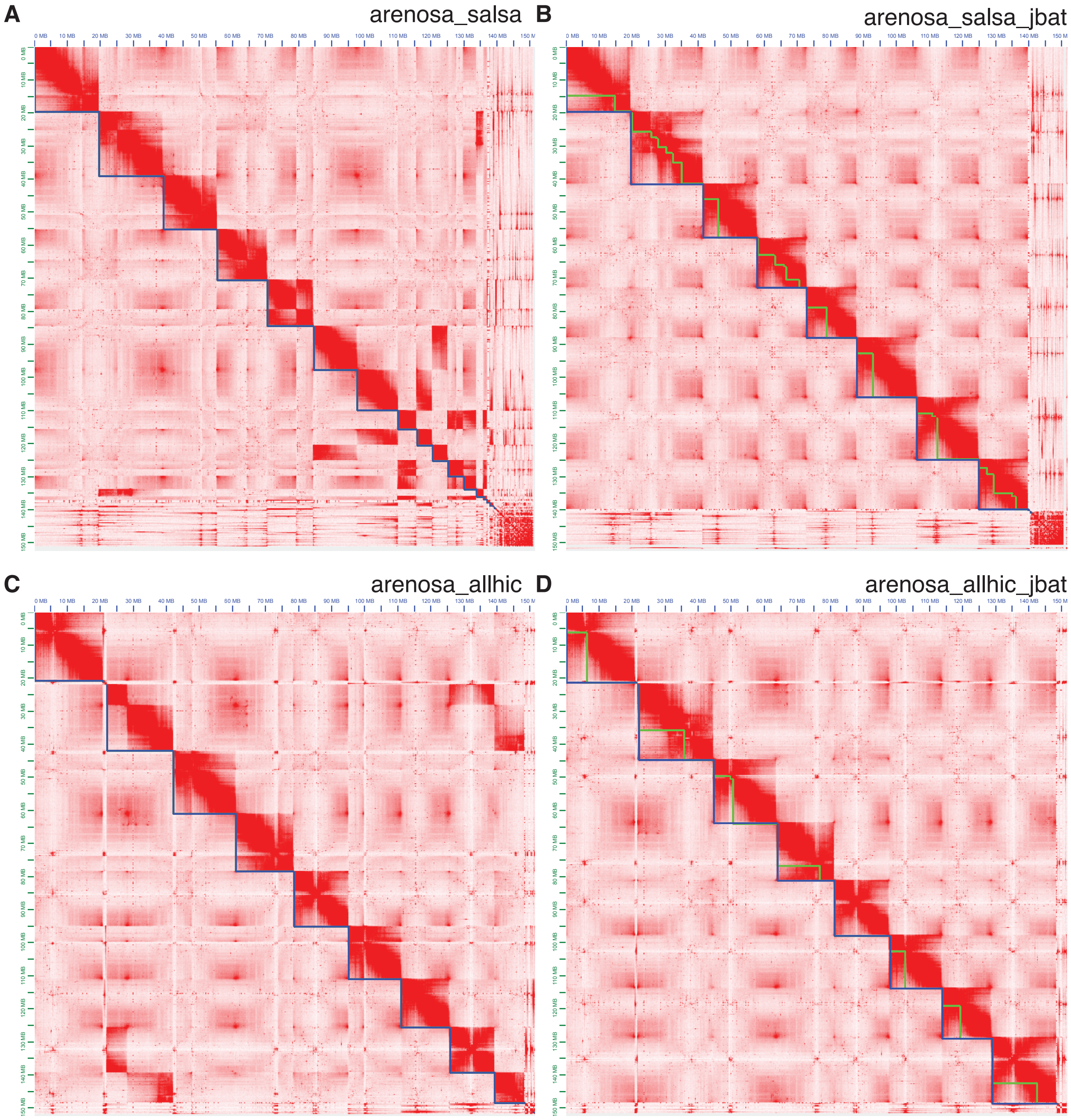


**Figure S6: *A. arenosa* Pusté Pole Hi-C contact maps.** Hi-C contact heat maps indicate the number of contacts between any given pair of loci in the assembly (red scale). Scaffolds are indicated by a blue line, and the green line documents the manual separation used for rearrangements. (A) Hi-C assembly heat map for *A. arenosa* Pusté Pole produced by the SALSA scaffolder. Strong signals far away from the diagonal were used for further scaffolding improvements. (B) The resulting contact heat map of the JBAT curated SALSA scaffolds. (C) Hi-C map of the *A. arenosa* Pusté Pole assembly scaffolded by the ALLHiC software. Problematic sequences are indicated by the low number of connections and stronger far-away signals. (D) Contact matrix after JBAT curation of miss joints and inversions. Green triangles indicate scaffold brakes. This assembly was selected and finalized by gap-filling with PBJelly (Figures S5 E and F).
