## Supplementary Figure S7: Genome alignments for "Structural evidence for MADS-box type I family expansion seen in new assemblies of *A. arenosa* and *A. lyrata*"

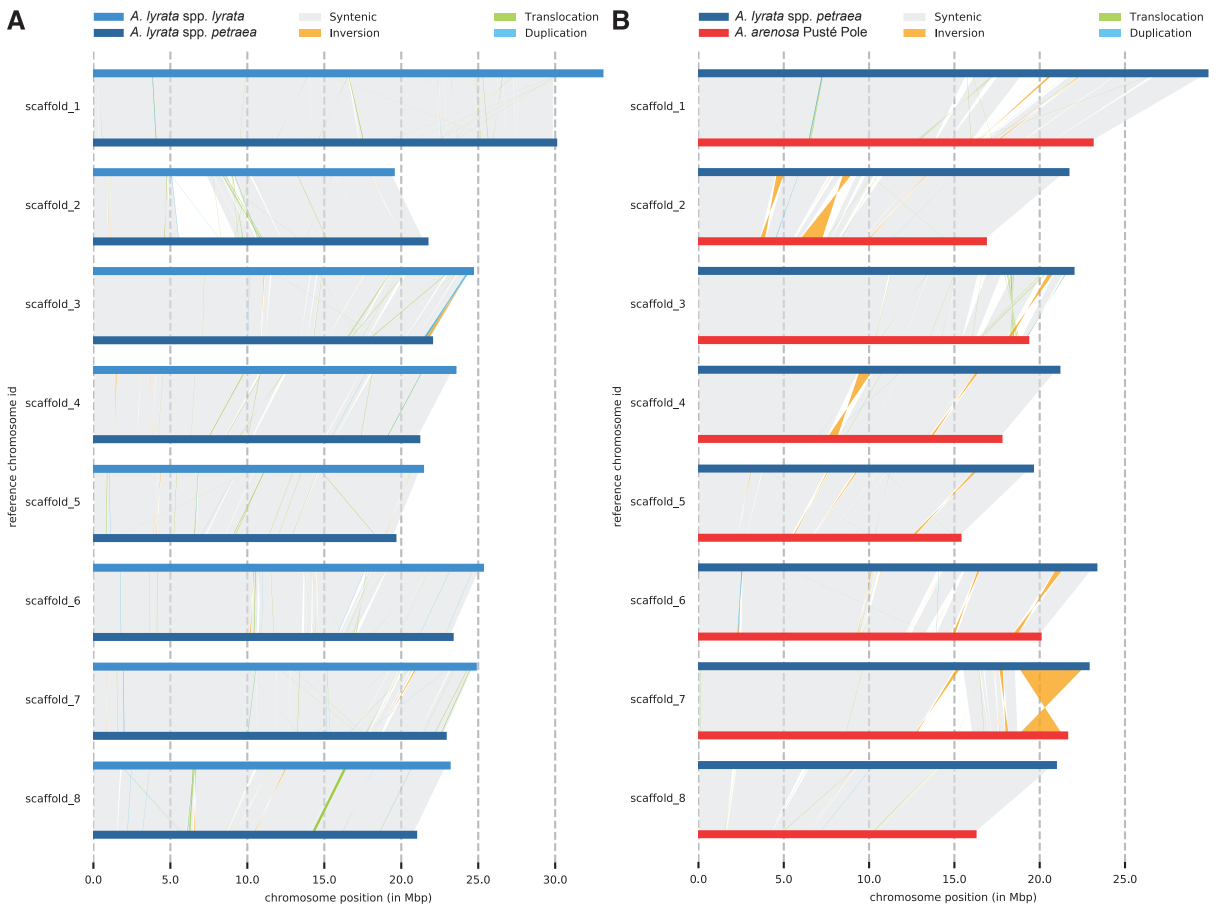


**Figure S7: Genome alignments indicate genomic rearrangements.** SyRI (Synteny and Rearrangement Identifier) was used to detect and classify structural differences between genomes. (A) Comparison of our *A. lyrata* ssp. *petraea* assembly (dark blue) to the *A. lyrata* ssp. *lyrata* reference (light blue). Syntenic regions are shown as gray blocks, while structural differences are grouped in translocations (green), inversions (orange), and duplications (blue). For visualization purposes, the relocations between non-homolog scaffolds have been excluded (e.g., for translocation between scaffolds one and two, see Figure 1E). (B) Synteny alignment between *A. lyrata* ssp. *petraea* (dark blue) and *A. arenosa* Pusté Pole (red). Structural differences are indicated between largely syntenic scaffolds.
