## Supplementary Figure S8: Extended MADS-box phylogeny for "Structural evidence for MADS-box type I family expansion seen in new assemblies of *A. arenosa* and *A. lyrata*"

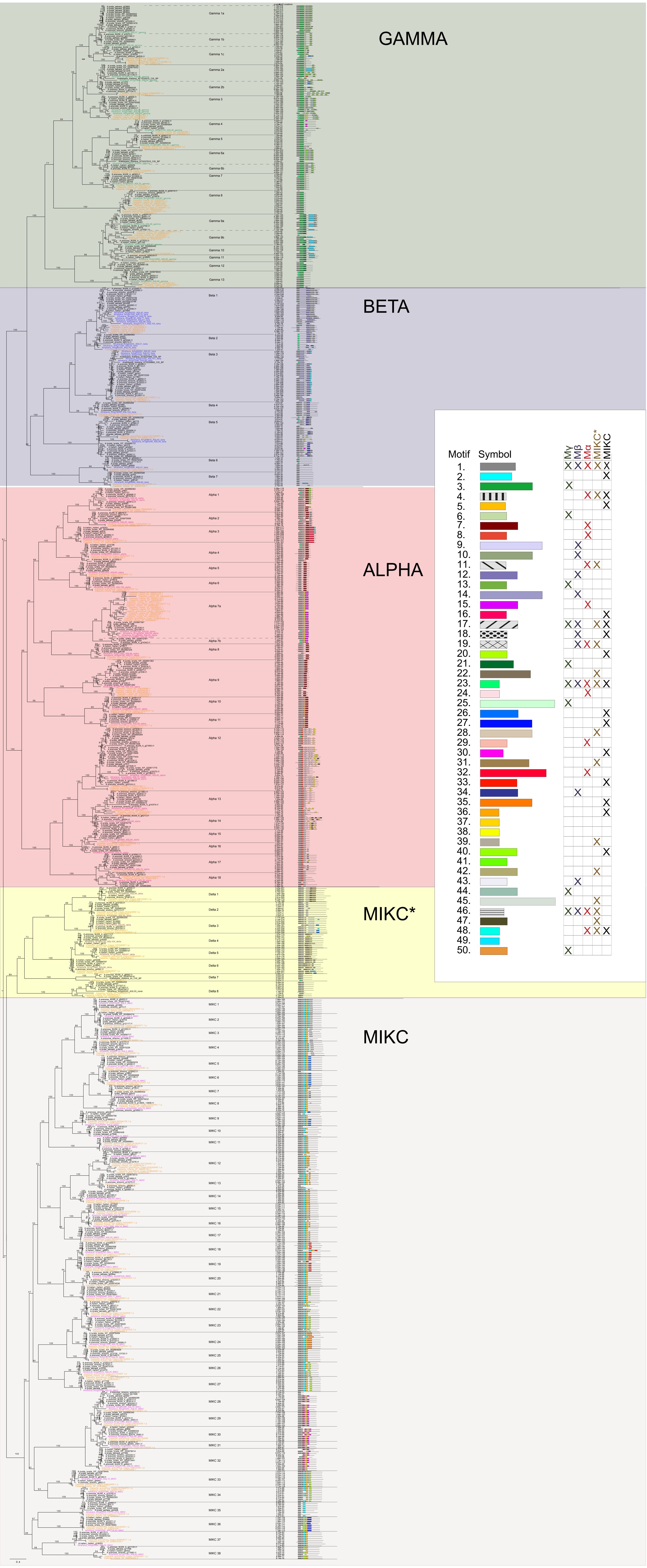


**Figure S8: Extended MADS-box phylogeny.** Phylogeny of all 841 identified MADS-box genes in *Arabidopsis* and *Capsella*. The groups correspond to previously published results with type I genes divided into three main groups (Mα, Mβ, Mγ), while type II genes fell into two monophyletic groups, MIKC and MIKC* [also referred to as Mδ] (Henschel et al., 2002; Parenicová et al., 2003; Arora et al., 2007; Thangavel and Nayar, 2018; Gramzow and Theißen, 2013; Qiu and Köhler, 2022). The groups are colored with a red box around Mα, blue around Mβ green around Mγ, yellow around MIKC*, and grey around MIKC. Sequences of *A. thaliana* are marked in blue, and the outgroup *Capsella* in orange. Monophyletic groups of *Arabidopsis* MADS-box genes, sharing a common ancestor with genes from the outgroup Capsella, are numbered starting from the top of the tree and delineated with solid lines. Additional gene duplications that occurred in the common ancestor of Arabidopsis, but after *Arabidopsis* and *Capsella* separated, are delineated with a dashed line and marked with an additional letter. For instance, “Mγ-1a” and “Mγ-1b” indicate that these two clades share a last common ancestor with *C. rubella* (i.e. “Mγ-1”), but have a gene duplication specific for *Arabidopsis* that occurred after the separation of *Arabidopsis* and *Capsella*. The right-hand column contains the result of the MEME analysis for each sequence. For the full result of the MEME analysis consult the material on GitHub. https://github.com/PaulGrini/Arabidopsis_assemblies
