## Supplementary Figure S9: MADS-box type II phylogeny for "Structural evidence for MADS-box type I family expansion seen in new assemblies of *A. arenosa* and *A. lyrata*"

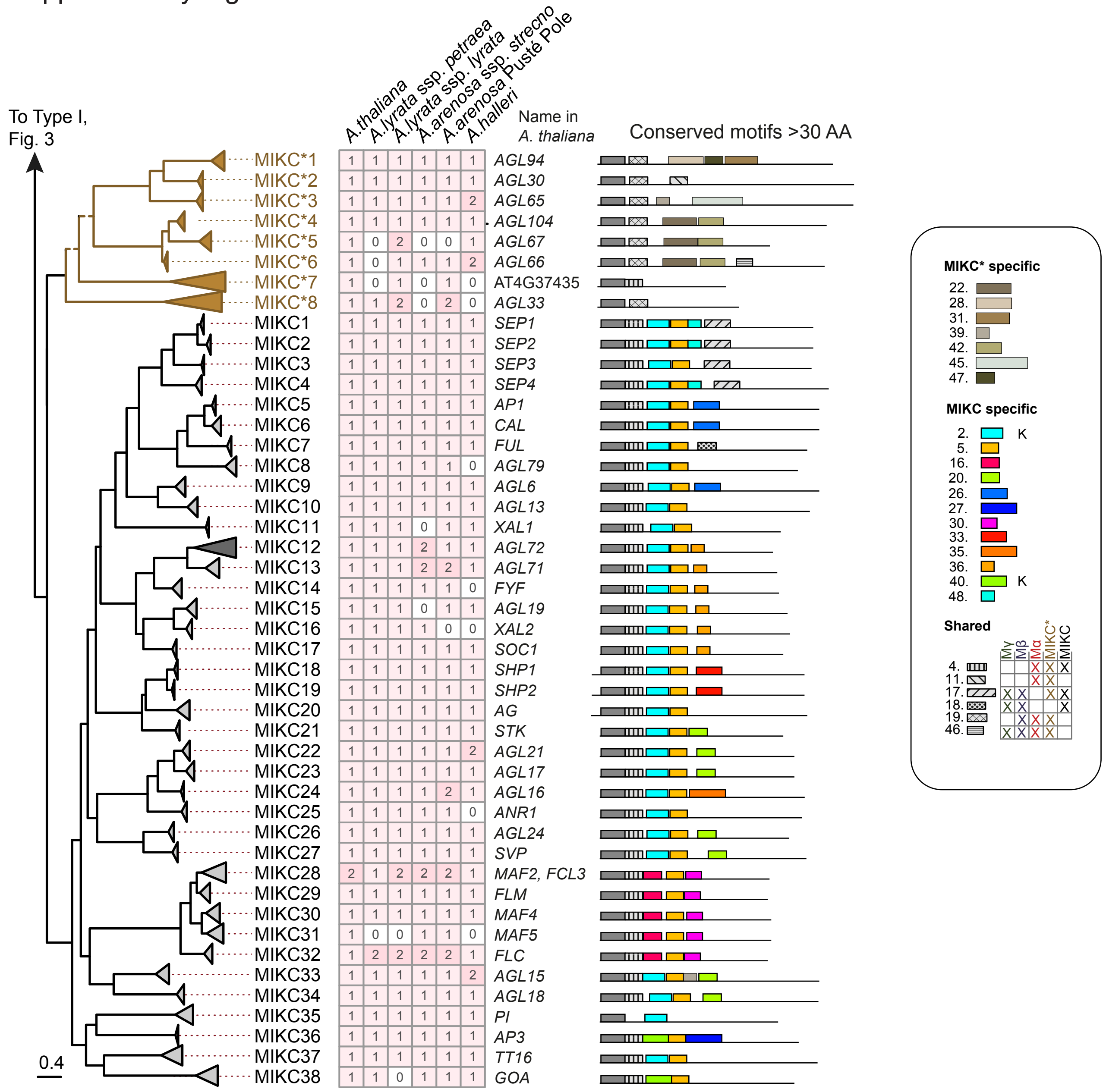


**Figure S9: Phylogenetic analysis of MADS-box type II genes in *Arabidopsis*.** The tree was derived by a maximum likelihood analysis of 275 identified MADS-box type I sequences from *A. thaliana, A. lyrata* ssp. *petraea, A. arenosa* Strecno, *A. arenosa* Pusté Pole*, and A. halleri* with two species of *Capsella* (89 sequences; not shown in the figure) used as outgroup. Solid branches represent bootstrap support > 85%, while branches with support values < 85% are dashed. The root of the tree is placed between type I and type II genes (the corresponding tree and heatmap for type I genes can be found in Figure 2). Triangles represent clades where branches are collapsed at the most recent gene duplication event in the last common ancestor of the genus *Arabidopsis*. The length of the triangles corresponds to the overall branch length of the collapsed clade, see the main text for naming schemes of clades. The heatmap shows the number of gene copies for each clade in the genomes of *A. thaliana, A. lyrata* ssp. *petraea, A. arenosa* Strecno*, A. arenosa* Pusté Pole*, and A. halleri.* The column next to the heatmap indicates the canonical AGL names of the genes in *A. thaliana* found in the corresponding clade; "none" means that a gene repre-senting the clade is not found in the *A. thaliana* genome, while "new" indicates that the gene does not have a given AGL name. The last column shows a simplified representation of the MEME motifs. A fully expanded phylogenetic tree with individual tip labels, support values for all branches, and the outgroup *Capsella* can be found in Figure S8. Results from the full MEME analysis can be found in Figure S8.
