## Supplementary Figure S10: PCA of MEME motifs for "Structural evidence for MADS-box type I family expansion seen in new assemblies of *A. arenosa* and *A. lyrata*"

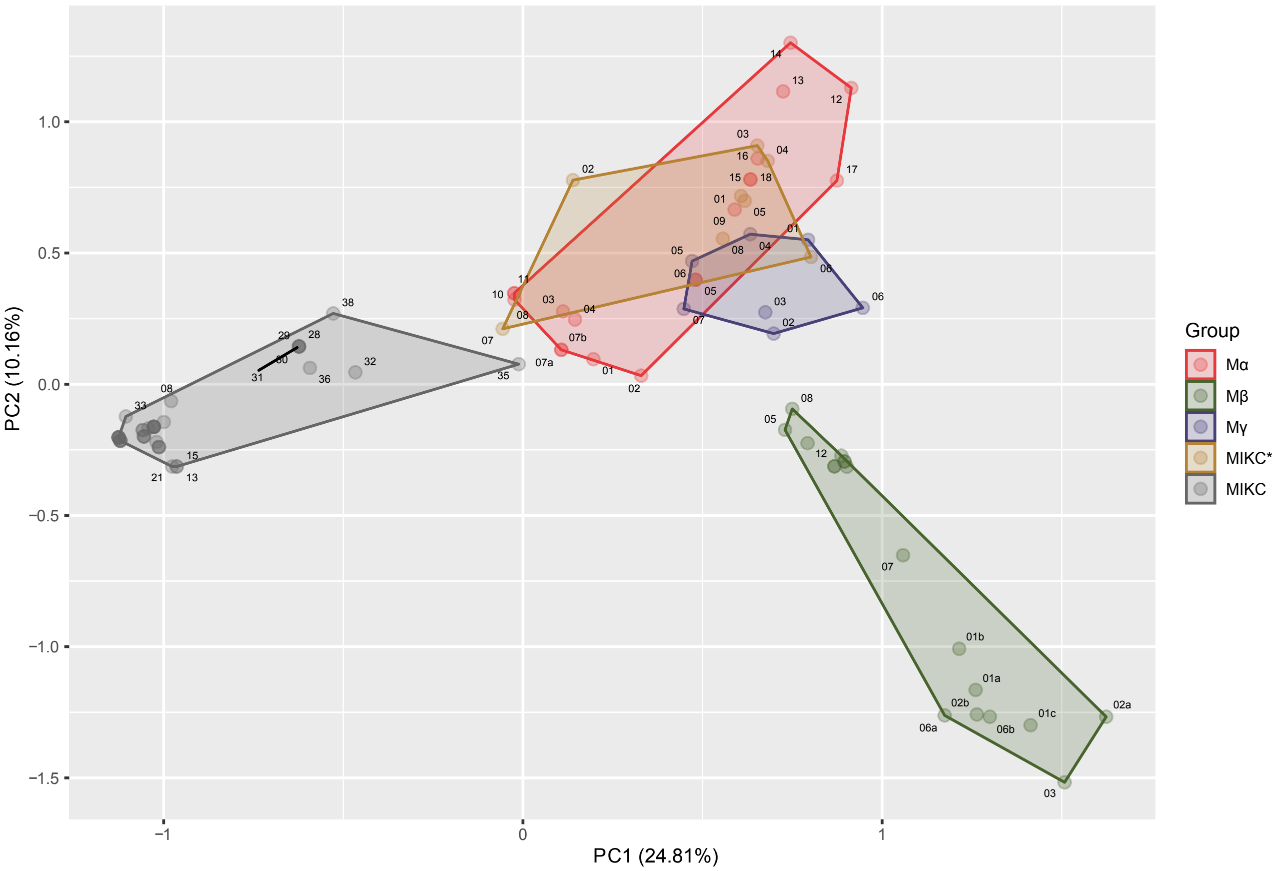


**Figure S10: PCA of the distribution of MEME motifs on MADS-box type I and type II genes**. The PCA was constructed from motifs identified by MEME and counted for each clade in the phylogeny. The first principal component axis captures 24.8% of the variation, and the second axis 10.2%. A polygon is drawn around each of the main groups with Mα in red, Mβ in blue, Mγ in green, MIKC* in yellow, and MIKC in grey.
