## Supplementary Figure S11: MADS-box type II expression during seed development for "Structural evidence for MADS-box type I family expansion seen in new assemblies of *A. arenosa* and *A. lyrata*"

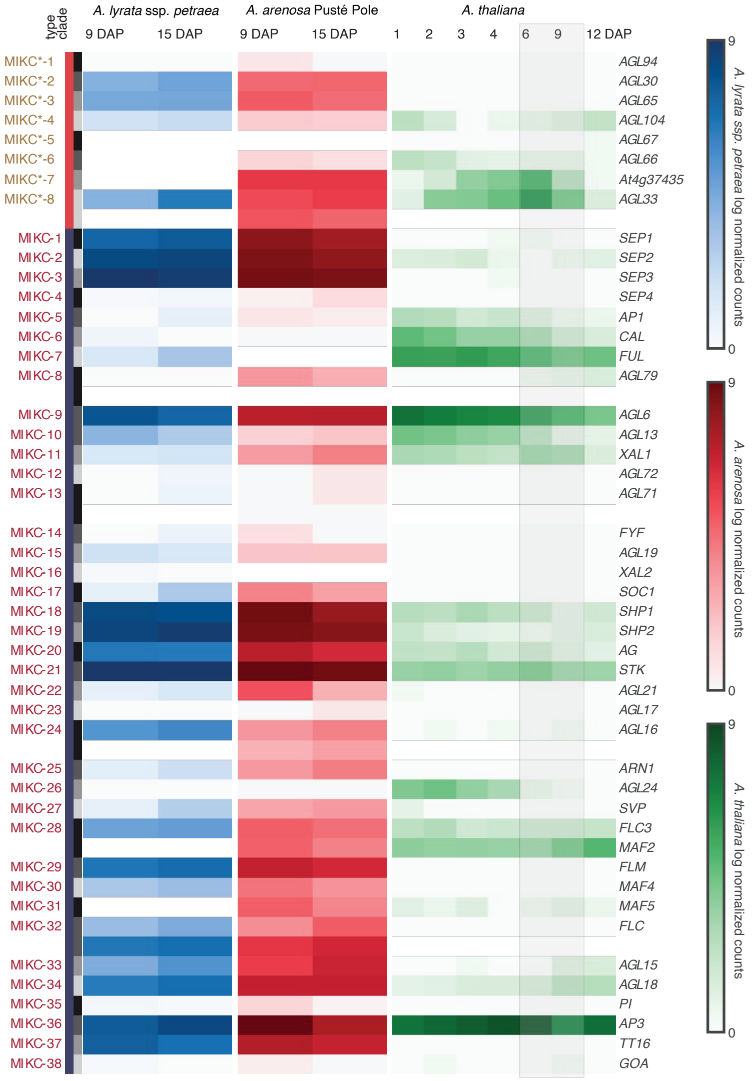


**Figure S11: MADS-box type II expression during seed development.** Gene expression profiles are displayed for all identified MADS-box type II genes and compared between *A. thaliana* (right column), *A. arenosa* Pusté Pole (middle column), and *A. lyrata* ssp. *petraea* (left column). The *A. thaliana* development time series (Bjerkan et al., 2020) serves as a reference. The endosperm cellularization in *A. thaliana* occurs between the 6 and 9 days after pollination (DAP). To adjust for the relatively slower development in *A. arenosa* Pusté Pole and *A. lyrata* ssp. *petraea*, corresponding stages before and after endosperm cellularization were sampled at 9 and 15 DAP, respectively. Ortholog genes are grouped and ordered and follow our MADS-box phylogeny (Figure 2). Sample normalized counts are shown with a base-2 logarithmic scale. The MADS-box type II seed expressions remain rather constant compared to the strong decline of type I expression around endosperm cellularization (Figure 4). However, expression differences can be seen between the *Arabidopsis* species. The *SEPALLATA (MIKC-1/2/3)* genes are strongly expressed in *A. lyrata* ssp. *petraea* and *A. arenosa* seeds while only weakly or absent in *A. thaliana*. In addition to their largely functional redundant role in flower development and ovary formation (Pelaz et al., 2001; Kaufmann et al., 2009), a crucial role in fruit development and ripening has been reported for *SEPALLATA* orthologs in tomato, strawberry, and apple (Ampomah-Dwamena et al., 2002; Seymour et al., 2011; Schaffer et al., 2013). Furthermore, the orthologs of *MIKC-15 (AGL19), MIKC-24 (AGL16), MIKC-26 (AGL24)* as well as the FLC clade *MIKC-28* to *MIKC-32* show strong expression differences reflecting the different regulations of flowering and strengthen of vernalization between the perennial *A. lyrata* ssp. *petraea* and *A. arenosa* plants compared to the annual *A. thaliana* (Schönrock et al., 2006; Alexandre and Hennig, 2008; Hu et al., 2014; Müller-Xing et al., 2022; Kemi et al., 2013; Soppe et al., 2021). We detect differing expressions of *MIKC**s between the species. *MIKC*-2* and *MIKC*-3* (AGL30 and AGL65) are expressed in *A. lyrata* ssp. *petraea* and *A. arenosa* seeds but are not detected in *A. thaliana*. On the other hand, *MIKC*-6* and *MIKC*-7* (*AT4G37435* and *AGL33*) show expression in *A. arenosa* and *A. thaliana* but not in *A. lyrata* ssp. *petraea* seeds. MIKC*s are known for their transcription activity in pollen (Verelst et al., 2007) but have also been detected in endosperm (Zhang et al., 2018). In addition, a high level of redundant heterodimers has been reported and double mutants reduce pollen fertility (Adamczyk and Fernandez, 2009).
