## Supplementary Figure S12: MAFFT alignments of genomic sequences for "Structural evidence for MADS-box type I family expansion seen in new assemblies of *A. arenosa* and *A. lyrata*"

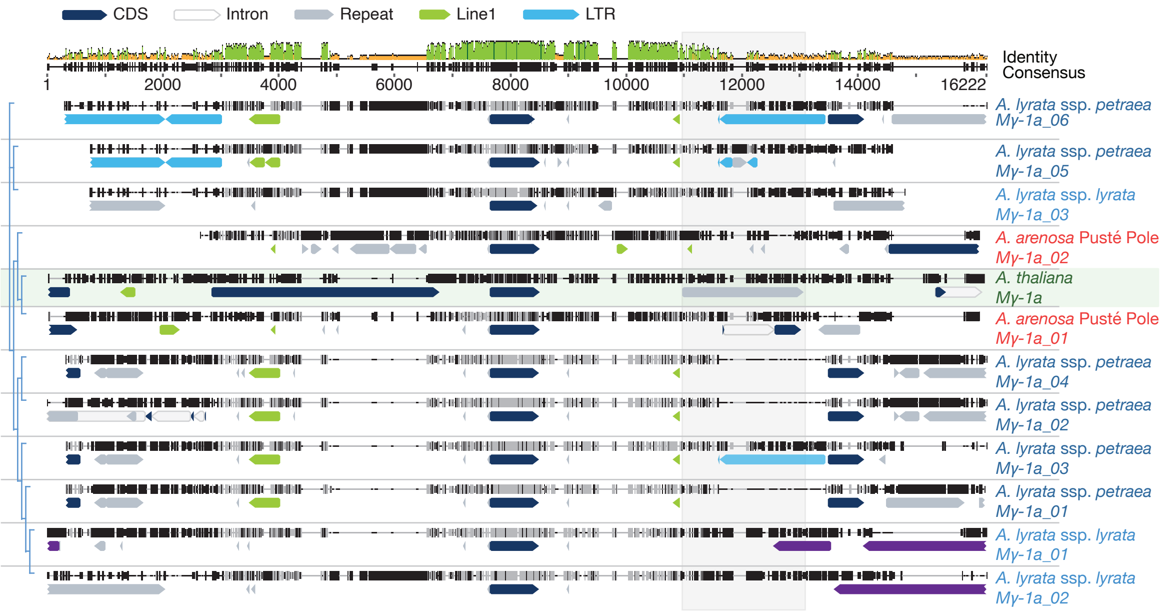


**Figure S12: MAFFT alignments of genomic sequences containing and surrounding Mγ-1**. Protein coding sequences (CDS), which are displayed as dark blue arrows of PHE1 orthologs from *A. thaliana* (green), *A. arenosa* (red), *A. lyrata ssp. lyrata* (light blue) and *A. lyrata* ssp*. petraea* (dark blue), are placed in the center. *A. thaliana* is highlighted by a light green background. On top, the consensus with identity score indicates a high similarity of the 3 prime regions downstream of the Mγ-1 loci. Around 2200 bp 3 prime of *PHE1* lies the repeat-rich region crucial for its parental-specific expression (inside the AT1TE79790 RC/Helitron, highlighted by a transparent gray box). Similar locations of Line 1 (light green arrows) and LTR transposons (light blue arrows) can be found next to the *Mγ-1* loci between species. Further repeats are marked as light gray arrows and lncRNA as violet arrows (XR_002328948.1 and XR_002334149.1).
